# Paf1 complex subunit Rtf1 stimulates H2B ubiquitylation by interacting with the highly conserved N-terminal helix of Rad6

**DOI:** 10.1101/2022.12.22.521652

**Authors:** Tasniem Fetian, Brendan M. McShane, Nicole L. Horan, Donya N. Shodja, Jason D. True, Amber L. Mosley, Karen M. Arndt

**Affiliations:** Department of Biological Sciences, University of Pittsburgh, Pittsburgh PA 15260; Molecular and Cellular Biology Graduate Program, University of Washington, Seattle WA 98195; Center for Developmental Biology and Regenerative Medicine, Seattle, Children’s Research Institute, Seattle WA 98101; Department of Biological Sciences, George Washington University, Washington, DC 20052; Department of Biochemistry and Molecular Biology, Indiana University School of Medicine, Indianapolis IN 46202; Department of Biology, Ball State University, Muncie, IN 47306

## Abstract

Histone modifications coupled to transcription elongation play important roles in regulating the accuracy and efficiency of gene expression. The mono-ubiquitylation of a conserved lysine in H2B (K123 in *Saccharomyces cerevisiae*; K120 in humans) occurs co-transcriptionally and is required for initiating a histone modification cascade on active genes. H2BK123 ubiquitylation (H2BK123ub) requires the RNA polymerase II (RNAPII)-associated Paf1 transcription elongation complex (Paf1C). Through its Histone Modification Domain (HMD), the Rtf1 subunit of Paf1C directly interacts with the ubiquitin conjugase Rad6, leading to the stimulation of H2BK123ub *in vivo* and *in vitro*. To understand the molecular mechanisms that target Rad6 to its histone substrate, we identified the site of interaction for the HMD on Rad6. Using *in vitro* crosslinking followed by mass spectrometry, we localized the primary contact surface for the HMD to the highly conserved N-terminal helix of Rad6. Using a combination of genetic and biochemical experiments, we identified separation-of-function mutations in *S. cerevisiae RAD6* that greatly impair H2BK123 ubiquitylation but not other Rad6 functions. Finally, by employing RNA-sequencing as a sensitive approach for comparing mutant phenotypes, we show that mutating either side of the proposed Rad6-HMD interface yields strikingly similar transcriptome profiles that extensively overlap with those of a mutant that lacks the site of ubiquitylation in H2B. Our results fit a model in which a specific interface between a transcription elongation factor and a ubiquitin conjugase guides substrate selection toward a highly conserved chromatin target during active gene expression.

**Significance Statement:** Transcription by RNAPII is tightly coordinated with mechanisms that control chromatin structure. Disruption of this interplay leads to deleterious effects on gene expression and genome architecture. Proteins that associate with RNAPII during transcription elongation play an important role in coupling histone modifications to active transcription. Paf1C, a conserved member of the RNAPII active elongation complex, is required for the ubiquitylation of histone H2B, a modification with effects on nucleosome stability and the methylation and acetylation state of chromatin. Here, we provide new insights into how a conserved domain in Paf1C, which we previously showed to be necessary and sufficient for Paf1C-mediated stimulation of H2B ubiquitylation, interacts with the ubiquitin conjugase for H2B thereby guiding its specificity.

## Introduction

Histone post-translational modifications (PTMs) are key to chromatin regulation during gene expression. The attachment of chemical groups or peptides to specific histone residues controls recruitment of regulatory proteins to nucleosomes and imposes structural changes to chromatin (1). The deposition of histone PTMs, like ubiquitylation, methylation, acetylation, and phosphorylation, typically involves a network of participating proteins. A prominent example is the mono-ubiquitylation of histone H2B on K123 in *S. cerevisiae* or K120 in mammals. The reversible ubiquitylation of H2B occurs co-transcriptionally and its level reflects the elongation rate of RNAPII (2–4). H2BK123ub is important for nucleosome stability during transcription elongation *in vivo*, likely in coordination with the histone chaperone FACT, and poses an energetic barrier to RNAPII *in vitro* (5–8). Through a crosstalk mechanism, H2BK123ub is required for the di- and tri-methylation of H3K4 and H3K79 via the Set1 and Dot1 methyltransferases, respectively (9–13). In turn, H3K4me2 and H3K4me3 recruit histone acetyltransferase and deacetylase complexes, further modulating chromatin accessibility (14–16).

The ubiquitin conjugase (E2) Rad6, the ubiquitin ligase (E3) Bre1, and the Bre1-associated protein Lge1 are required for H2BK123ub in *S. cerevisiae*, and homologs of Rad6 and Bre1 function analogously in other eukaryotes, including humans (17–21). As an E2, Rad6 coordinates with different E3 proteins to ubiquitylate substrates within diverse biochemical pathways. This is reflected in the numerous mutant phenotypes associated with *rad6* mutants. For example, in yeast, null alleles of *rad6* affect silencing of transcription near telomeres (18, 22), post-replication repair of DNA damage (23), protein degradation dependent on the N-end rule pathway (24, 25), response to oxidative stress (26), and Ty1 element transposition (27, 28).

During transcription elongation, RNAPII interacts with regulatory proteins that co-transcriptionally modulate its activity and/or the local chromatin environment. The evolutionarily conserved, multifunctional Paf1 complex (Paf1C), comprised of five subunits in yeast (Paf1, Rtf1, Ctr9, Cdc73, and Leo1), is a core component of the RNAPII active elongation complex that facilitates transcription elongation across eukaryotic genomes (29). In addition, Paf1C is important for coordinating processes coupled to elongation, including transcription termination and RNA 3’-end formation. Importantly, Paf1C also promotes the deposition of several co-transcriptional histone modifications (29). In yeast and other eukaryotes, Paf1C is required for H2B mono-ubiquitylation (30–33) in a manner dependent on a small domain within the Rtf1 subunit, which we termed the Histone Modification Domain (HMD) (32, 34–36). Demonstrating its pivotal role in stimulating H2BK123ub, heterologous expression of the HMD is sufficient to restore H2BK123ub in a yeast strain lacking all five subunits of Paf1C (33). The ability of a recombinant HMD to stimulate H2BK123ub in a reconstituted, transcription-free system argues that Paf1C functions directly in stimulating the ubiquitylation of H2B by Rad6-Bre1 (33). In addition to Rtf1, recent work has shown that other Paf1C subunits contribute to efficient H2B ubiquitylation (33, 37).

The molecular mechanisms underlying how Rad6 and Bre1 specifically target H2B and the role of Paf1C in this process are not fully understood. We previously demonstrated that expression of the HMD outside the context of intact Paf1C restores H2BK123ub levels but leads to mislocalization of the modification, arguing that Paf1C helps control H2BK123ub patterning via tethering the HMD to RNAPII (33). Using site-specific crosslinking, we showed that the HMD directly interacts with Rad6 *in vivo* and identified the region within the HMD that interacts with Rad6 (33). However, the interaction region within Rad6 remained uncharacterized. The significance of this question is underscored by the multiple functions of Rad6, which, as only one of eleven E2 proteins in budding yeast (38), is targeted to its appropriate substrates through dynamic interactions with other proteins. Here, through site-specific crosslinking and separation-of-function mutations, we demonstrate that the N-terminal alpha helix of Rad6 interacts with the HMD and is critical for HMD-mediated stimulation of H2BK123ub. Our findings suggest a highly specific interface between Paf1C and Rad6 that helps direct Rad6 to its nucleosomal substrate.

## Results

### The Rtf1 HMD interacts with the N-terminal alpha helix of Rad6 *in vitro*

Given the dynamic nature of transcription elongation and associated histone modifications, we used a site-specific *in vitro* crosslinking strategy to identify the region of Rad6 that interacts with the Rtf1 HMD. Briefly, an engineered tRNA/aminoacyl-tRNA synthetase system and amber codon suppression in *E. coli* were used to express an HMD_74-184_ protein (residues 74-184 of Rtf1) with the photoreactive amino acid analogue *p*-benzoyl-L-phenylalanine (BPA) at amino acid 108, replacing a natural phenylalanine at that position (39). Upon photoactivation, BPA crosslinks to residues within a 10 Å distance (40). We previously showed that Rtf1-F108BPA crosslinked to Rad6 *in vivo* and substitution of F108 with alanine greatly diminished cellular levels of H2BK123ub (33). We purified the HMD_74-184-F108BPA_ protein (Fig. S1A) and showed that it was able to stimulate the levels of H2BK123ub similar to wild-type (WT) HMD in a reconstituted reaction containing recombinant yeast Rad6 (yRad6), yBre1, human E1 (hE1), ubiquitin, and *Xenopus laevis* mononucleosomes (Fig. S1B). The HMD_74-184_-F108BPA protein was incubated with recombinant V5-tagged yRad6 alone (Fig. 1A) or in the context of a complete H2B ubiquitylation reaction +/− ATP (Fig. 1B and Fig. S1C), and the reaction mixtures were exposed to long-wave UV light to activate crosslinking by BPA. In the presence or absence of other factors or catalysis, the HMD_74-184-F108BPA_ protein formed a crosslink with Rad6, as detected by western blot analysis when probing for either the HMD (Fig. S1C) or Rad6 (Fig. 1A-B). Liquid chromatography tandem mass spectrometry (LC-MS/MS) analysis of the V5-Rad6~HMD photo-crosslinked product, which was gel purified from a reaction containing only V5-Rad6 and HMD_74-184-F108BPA_, identified residues in Rad6 that were directly crosslinked to BPA. The highest number of peptide spectral matches mapped to amino acids 9-12 in the highly conserved N-terminal alpha helix of Rad6 (Fig. 1C-D), although we also detected a small number of crosslinks to a C-terminal region of Rad6 and a large number to the N-terminal V5 tag (Table S1). In the absence of additional factors, the HMD-Rad6 interface might be dynamic, explaining the appearance of crosslinks to multiple residues. These data demonstrate that Rad6 directly interacts with the Rtf1 HMD *in vitro* through its highly conserved N-terminal helix and provide a foundation for genetic dissection of this interaction.

**Figure 1:**
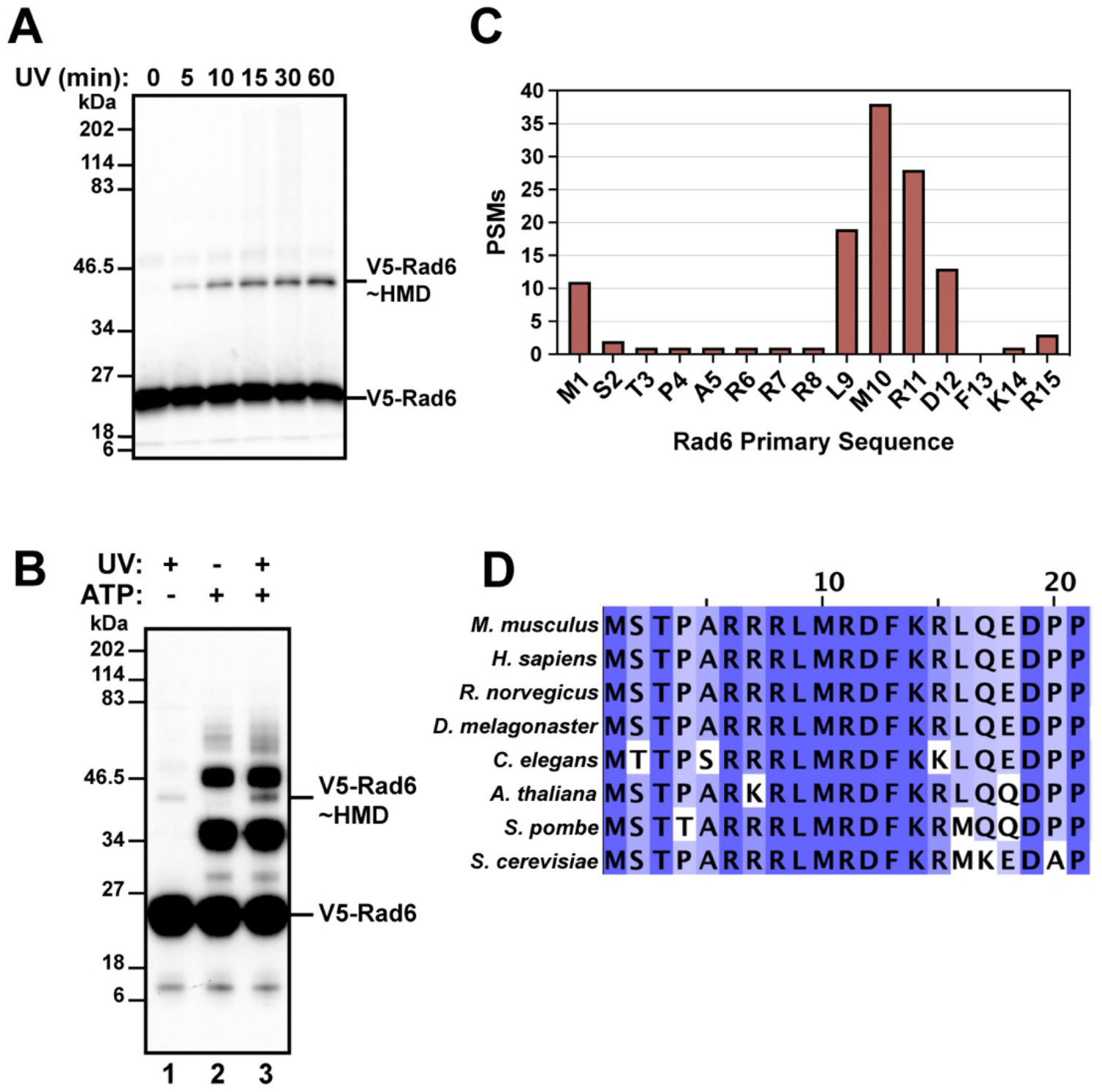
*In vitro* site-specific crosslinking identifies a conserved HMD-interacting region within the Rad6 N-terminus. (A) Recombinant V5-Rad6 and HMD_74-184-F108BPA_ were mixed and exposed to UV light (365 nm) for the indicated periods of time. Crosslinking was monitored by western analysis using anti-V5 antibody. (B) Western blot probed with anti-V5 antibody to detect crosslinked products in a complete H2B ubiquitylation reaction containing V5-Rad6, HMD_74-184-F108BPA_, yBre1, ubiquitin, hE1, and *X. laevis* nucleosomes with or without ATP. Reactions were exposed to UV light as indicated. Lane 1 lacked ATP and contained apyrase. The ATP-dependent high molecular weight V5-reactive species are likely ubiquitylated forms of V5-Rad6. (C) Mass spectrometry analysis of crosslinked V5-Rad6~HMD_74-184-F108BPA_, quantified as peptide spectrum matches (PSMs), identifies residues in the N-terminal helix of Rad6 that crosslinked to the HMD *in vitro*. (D) Multiple sequence alignment of the Rad6 N-terminal region. A darker color indicates higher conservation across species. This panel was generated using Jalview (68).

### Alanine scanning mutagenesis identifies Rad6 residues specifically required for H2BK123ub *in vivo*

Guided by our *in vitro* crosslinking data, we subjected the N-terminal alpha helix of Rad6 to alanine scanning mutagenesis to identify amino acids required for H2BK123ub in yeast. Plasmids expressing WT or mutant derivatives of Rad6 were transformed into a *rad6*Δ strain and histone modifications were assessed by western blotting. Alanine substitutions at positions 10 and 11 of Rad6, which had been identified as HMD_74-184-F108BPA_ crosslinking sites (Fig. 1C), and also at positions 2, 3, 6, and 8 severely reduced global H2BK123ub levels (Fig. 2A and Fig. S2A). Levels of H2BK123ub-dependent modifications, namely H3K4me3 and H3K79me2/3, were also reduced in these mutants (Fig. 2A). Substitutions at positions 12 and 13 of Rad6 caused a more modest reduction in H2BK123ub levels (Fig. 2A and Fig. S2A). In all *rad6* substitution mutants, Rad6, Bre1, and Rtf1 levels were similar to WT levels, arguing that the loss of H2BK123ub is not due to decreased levels of these important H2B ubiquitylation factors (Fig. 2A, Fig. S2B).

**Figure 2:**
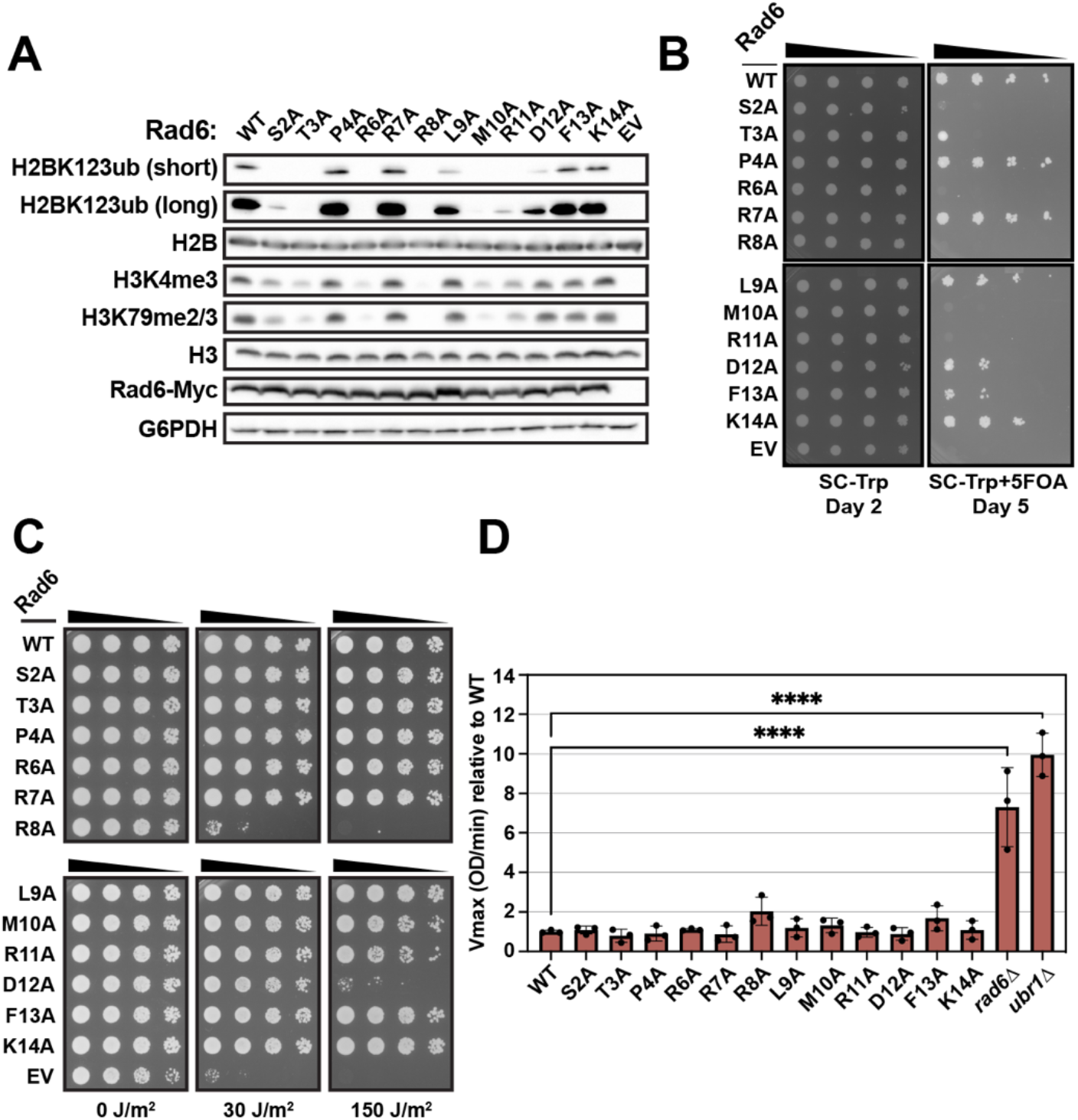
Alanine scanning mutagenesis of Rad6 N-terminal helix identifies residues required for H2B ubiquitylation. (A) A *rad6*Δ strain (KY2045) was transformed with plasmids expressing WT *RAD6* or *rad6* mutant alleles, and extracts of transformants were analyzed by western blotting with antibodies against the indicated proteins. Short and long exposures of the same H2BK123ub blot are shown. EV = empty vector. G6PDH serves as a loading control. (B) A *rad6*Δ strain (KY3391) bearing a telomeric silencing reporter (*TELVR::URA3*) was transformed with WT or mutant *rad6* plasmids, and transformants were spotted in a five-fold serial dilution series onto SC-Trp+5FOA or growth control media (SC-Trp). (C) Five-fold serial dilution spot assay of KY2045 transformants followed by exposure to indicated doses of UV irradiation. Representative images show growth on day two relative to the time of plating. (D) Function of alanine-substituted *rad6* mutants in protein degradation via the N-end rule pathway was assayed using a β-galactosidase reporter with a destabilizing residue (Arg) at the N-terminus (41). Relative V_max_ values for β-galactosidase were calculated with reference to the average V_max_ from the WT measurements. All strains were assayed in biological triplicate and technical duplicate. Transformants of strain KY3392 contained two plasmids: the Arg-β-galactosidase reporter (pKB1526) and a WT or mutant *rad6* plasmid. The *ubr1Δ* control strain was constructed by transforming the WT *RAD6* plasmid (pKB1167) into a *rad6Δ ubr1Δ strain* (KY3389). Mean and SD are shown. Significant differences compared to the WT are indicated by asterisks based on an unpaired one-way ANOVA with Dunnett’s post-hoc test (****p<0.0001).

To determine if the *rad6* mutants exhibit phenotypes associated with the loss of H2BK123ub specifically or if they are more broadly impaired in Rad6 functions, we performed a set of phenotypic tests. Consistent with the requirement for H2BK123ub in the silencing of genes positioned near telomeres (9), the *rad6* mutants that have the greatest defects in H2BK123ub (substitutions S2A, T3A, R6A, R8A, M10A and R11A) are all defective in silencing a *URA3* reporter gene positioned near a telomere (Fig. 2B). These mutants are strongly sensitive to media containing 5FOA, a compound that is toxic to *URA3*-expressing yeast cells, while mutants with WT levels of H2BK123ub (P4A, R7A, and K14A) grow normally on this media (Fig. 2B). Mutants with intermediate defects in H2BK123ub (D12A and F13A) show only a modest reduction in growth (Fig. 2B).

In contrast to the strong correlation between telomeric silencing and H2BK123ub defects, only one *rad6* substitution mutant, *rad6-R8A*, was strongly sensitive to UV irradiation, a phenotype that reflects the ability of Rad6 to modify PCNA and promote DNA damage repair via translesion synthesis (Fig. 2C) (23). One other mutant, *rad6-D12A*, was sensitive to the highest dose of UV exposure, but none of the other H2BK123ub-defective *rad6* mutants shared this property. Another well-established process dependent on Rad6 is proteolysis of substrates through the N-end rule pathway (24, 25). To test for defects in N-end rule function in the *rad6* mutants, we used a well-characterized plasmid-based reporter expressing an unstable substrate: β-galactosidase with a strongly destabilizing amino acid, Arg, at the N-terminal position (41). Degradation of this substrate is dependent on Rad6 and the E3 Ubr1 (24, 25, 42) (Fig. 2D). In contrast to *rad6*Δ and *ubr1*Δ strains and when compared to WT, the single alanine substitutions caused little if any effect on the levels of the N-end rule substrate, as measured by β-galactosidase assays in extracts from the mutants (Fig. 2D). Finally, a recent study demonstrated a role for Rad6 in the cellular response to oxidative stress through ubiquitylation of ribosomal proteins (43). Exposure of yeast cells to hydrogen peroxide revealed strong sensitivity for the *rad6*Δ mutant, modest sensitivity for the *rad6-R8A* mutant, and WT levels of resistance for the other mutants (Fig. S2C). As summarized in Fig. S2D, our genetic analysis of the Rad6 N-terminal helix has identified mutations that impair H2BK123ub and processes requiring this mark but apparently do not affect three other Rad6-dependent pathways.

### Separation-of-function mutations identify Rad6 residues required for stimulation by the HMD

The lack of H2BK123ub in the *rad6* mutants could arise through multiple mechanisms, including disrupted binding of Rad6 to the HMD, Bre1, nucleosomes, or other factors. Therefore, to specifically test the ability of the Rad6 substitution mutants to respond to stimulation by the HMD, we turned to the *in vitro* H2B ubiquitylation assay assembled with recombinant factors. For these experiments, we purified WT and mutant Rad6 proteins (Fig. S3A); each of the nine mutants selected was defective for H2BK123ub *in vivo* (Fig. 2A and Fig. S2A).

In agreement with our prior work (33), recombinant WT HMD_74-184_ can stimulate the level of Bre1-dependent H2BK123ub in the reconstituted reaction approximately three-fold (Fig. 3A-B, compare hatched and filled bars in B). The Rad6 mutants fell into three groups. In the first group, the R8A and R11A mutants are strongly defective for H2BK123ub *in vitro* even in the basal condition, defined as in the presence of Bre1 but not HMD (Fig. 3A-B; Fig. S3B). For R8A, this strong defect correlates with phenotypes in addition to the loss of H2BK123ub and telomeric silencing, including UV sensitivity on par with the *rad6*Δ mutant (Fig. 2C) and moderate sensitivity to hydrogen peroxide (Fig. S2C). Within the structure of Rad6, R8 is engaged in a network of interactions that likely contribute to the structural fold (44). The behavior of the R11A mutant appears more specific to H2BK123ub and a lack of response to Bre1 (Fig. S3B), as this mutant confers a strong H2BK123ub defect yet supports WT levels of N-end rule activity and tolerance to UV and oxidative stress (Fig. 2 and Fig. S2). In the second group, mutants S2A and T3A surprisingly behaved like WT Rad6 *in vitro* despite their severe effect on H2BK123ub *in vivo* (Fig. 2A and Fig. S2B). This suggests that Rad6 residues S2 and T3 are required for a step in H2B ubiquitylation that manifests *in vivo*, possibly for the response to other factors like Lge1 (20, 45). In the third group, the Rad6 R6A, L9A, M10A, D12A and F13A mutant proteins all show reduced ability to be stimulated by the HMD (Fig. 3B, solid bars) yet support near WT or even greater (D12A) levels of H2BK123ub in the absence of the HMD (Fig. 3B, hatched bars).

**Figure 3:**
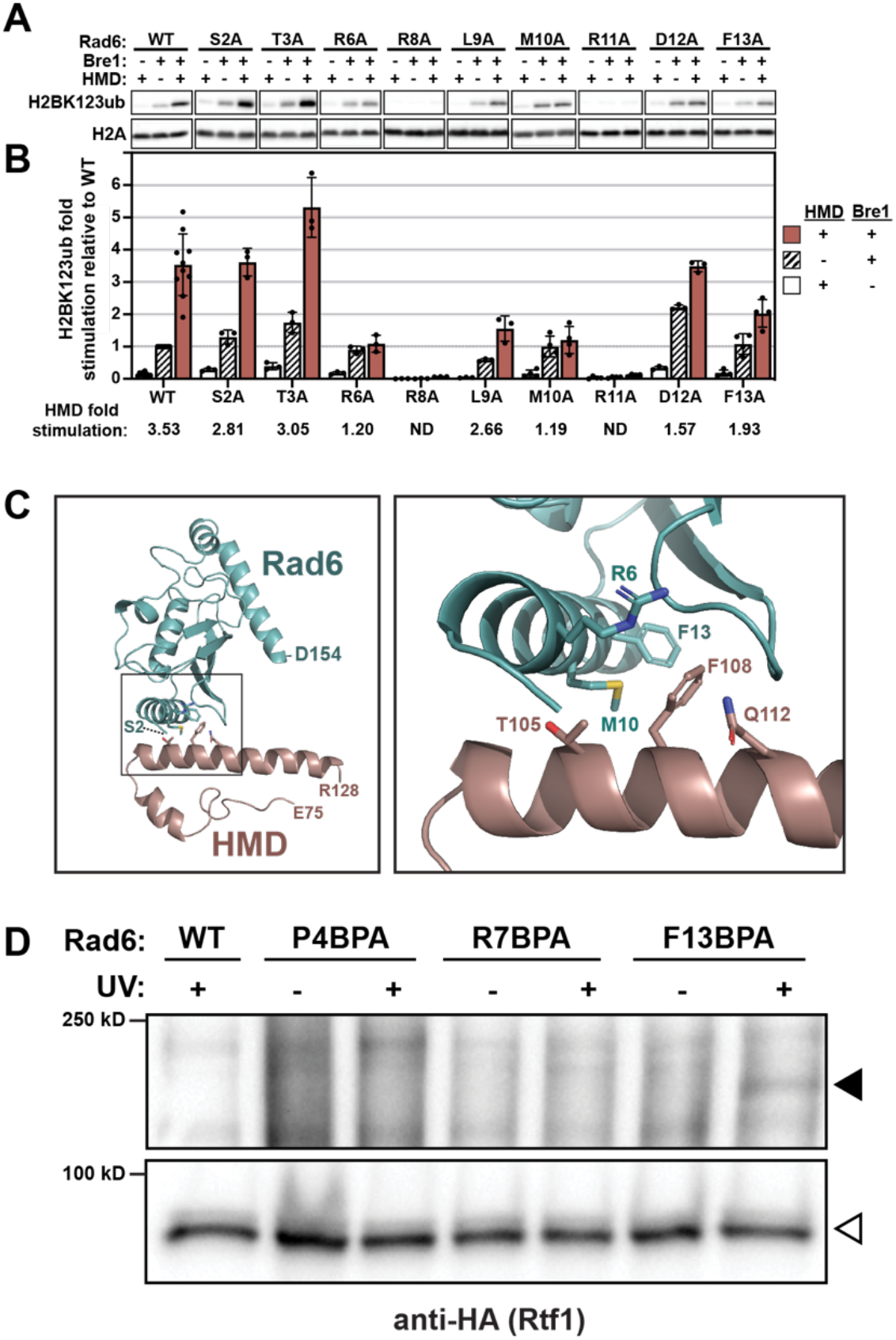
Specific residues in the Rad6 N-terminus are required for stimulation of H2B K123 ubiquitylation by the HMD *in vitro*. (A) Representative western blots show the extent of H2B K123 ubiquitylation in the absence of either HMD_74-184_ or yBre1 or in the presence of both. H2A serves as the loading control. (B) Quantification of western blots shown in A. Unfilled (white) bars represent reactions containing the HMD but lacking Bre1. Hatched bars represent reactions containing Bre1 but lacking the HMD. Red solid bars represent reactions containing both Bre1 and the HMD. H2BK123ub fold stimulation in all lanes was calculated relative to H2BK123ub-normalized signal from WT Rad6 in the presence of Bre1 and absence of the HMD (WT middle lane) (see SI Methods). Points show individual measurements (minimum of three technical replicates). Mean and SD are shown. Below the graph, H2BK123ub-normalized signals in both Bre1-containing reactions were used to calculate the HMD fold stimulation per mutant as the ratio between H2BK123ub in the presence and absence of the HMD. ND = not determined due to very low signals. (C) Crosslinking-instructed model of the Rad6-HMD interface using crystal structures of Rad6 (PDB:1AYZ) and Rtf1 HMD (PDB:5E8B). Docking was done using ClusPro 2.0. The inset in the left panel is expanded on the right. (D) *In vivo* BPA crosslinking followed by western analysis with antibody against the 3xHA tag on Rtf1 shows the appearance of a UV-dependent crosslinked species when Rad6-F13 is substituted with BPA in yeast. A *rad6*Δ strain (KY2798) was transformed with 2μ plasmids expressing *rad6* amber codon mutations, allowing BPA incorporation at the indicated locations in Rad6. Black arrowhead denotes the Rad6~3xHA-Rtf1 crosslinked product. White arrowhead indicates levels of un-crosslinked 3xHA-Rtf1.

With respect to fold stimulation by the HMD, the R6A and M10A mutants are the most strongly affected. Interestingly, M10 was the primary site of crosslinking between the HMD74-184-F108BPA protein and Rad6 *in vitro* (Fig. 1C).

Using published structures (33, 44) and instructed by our *in vitro* BPA crosslinking results, we generated a model of the putative Rad6-HMD interface using ClusPro 2.0 (Fig. 3C) (see SI Methods). This model positions R6 and M10 of Rad6 near HMD residues that crosslink to Rad6 *in vivo* (Rtf1 T105, F108, Q112) (33) and *in vitro* (Rtf1 F108; Fig. 1). In addition, we used Alphafold2-multimer (46, 47) to generate a model of the Rad6-HMD interaction independent of experimental constraints (Fig. S3C). Remarkably, the Alphafold2-multimer prediction places the Rad6 M10 residue in close proximity to Rtf1 F108 and other residues that crosslink to Rad6 *in vivo* (33). However, the two models differ in the orientation of Rad6 relative to the HMD. To begin testing these models, we incorporated BPA into Rad6, replacing amino acids P4, R7, or F13 via amber codon suppression in yeast (48). Alanine substitutions of these amino acids caused minor, if any, effects on H2BK123ub *in vivo* (Fig. 2A and S2A). Yeast transformants harboring the tRNA/aminoacyl-tRNA synthetase plasmid for BPA incorporation and a plasmid containing either WT *RAD6* or an amber codon derivative of *RAD6* were exposed to long-wave UV light. Consistent with the *in silico* models, Rad6-F13BPA reproducibly crosslinked to 3xHA-Rtf1, as assessed by western blotting (Fig. 3D). These results independently support our *in vitro* crosslinking, biochemical, and genetic data.

### Targeted *rad6* mutations specifically disrupt HMD-dependent H2BK123 ubiquitylation

Based on our molecular models and analysis of single substitution mutants, we hypothesized that Rad6 amino acids R6 and M10 are particularly important for the Rad6-HMD interaction. To uncover the effects of *rad6-R6A* and *rad6-M10A* on H2B ubiquitylation *in vivo* with greater sensitivity, we deleted *UBP8* and *UBP10*, the genes encoding the deubiquitylases that target H2BK123ub (3). Unlike results observed in the presence of *UBP8* and *UBP10* (Fig. 4A, left), deletion of these genes revealed residual H2BK123ub and higher levels of H3K4me3 and H3K79me2/3 in the *rad6-R6A, rad6-R8A* and *rad6-M10A* mutants indicating that they are hypomorphic for H2BK123ub loss (Fig. 4A, right).

**Figure 4:**
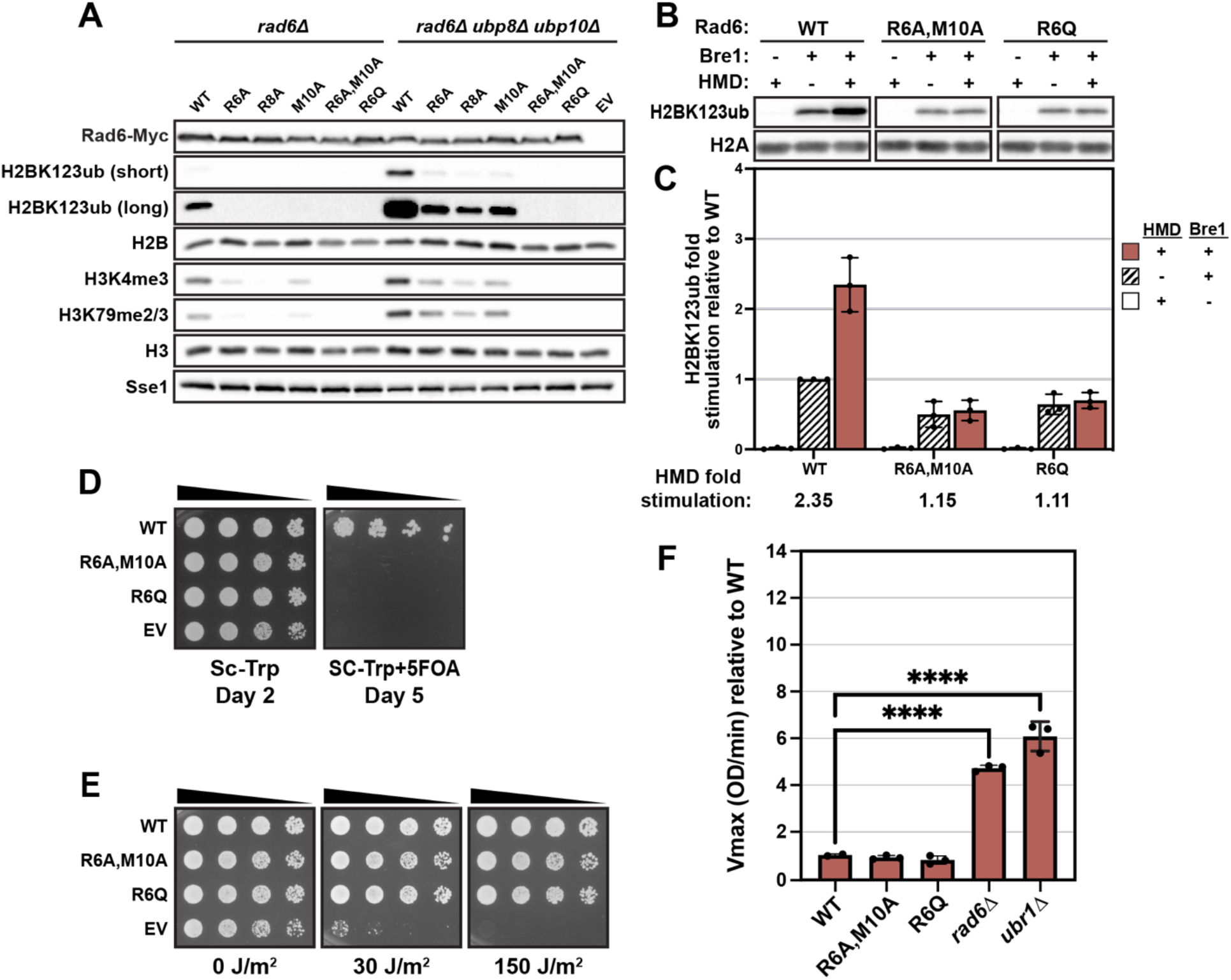
*rad6-R6Q* and *rad6-R6A, M10A* mutants are severely and specifically defective in the H2BK123ub pathway. (A) Either *rad6*Δ (KY2045) or *rad6*Δ *ubp8*Δ *ubp10*Δ (KY4010) strains were transformed with WT or *rad6* mutant plasmids followed by western analysis. Sse1 serves as a loading control. Short and long exposures of the same H2BK123ub blot are shown. EV = empty vector. (B, C) *In vitro* H2B ubiquitylation assay and associated quantitation (as in Fig. 3A-B) to assess the basal activity and response of Rad6-R6A, M10A and Rad6-R6Q proteins to HMD stimulation. (D) Five-fold serial dilution spot assay of *rad6*Δ *TELVR::URA3* (KY3391) transformants similar to Fig. 2B. (E) Five-fold serial dilution spot assay of *rad6*Δ (KY2045) transformants on SC-Trp medium followed by exposure to indicated doses of UV irradiation. Representative images show growth on day two relative to the time of plating. (F) Protein degradation via the N-end rule pathway was assayed as in Fig. 2D except that WT was measured in biological duplicate with one assayed in technical quadruplicate and the other in duplicate. Mean and SD are shown. Significant differences compared to the WT are indicated by asterisks based on an unpaired one-way ANOVA with Dunnett’s post-hoc test (****p<0.0001).

To generate *rad6* mutants more severely compromised for HMD-mediated stimulation of H2BK123ub, we constructed a *rad6-R6A, M10A* double mutant and also a *rad6-R6Q* mutant. The latter was chosen based on surveying human cancer databases (COSMIC, ClinVar, and NCBI dbSNP) to identify recurring missense mutations in the N-terminal region of either UBE2A or UBE2B, the human homologs of yeast Rad6. Strikingly, the *rad6-R6Q* and *rad6-R6A, M10A* mutants lack detectable H2BK123ub, H3K4me3 and H3K79me2/3 either in the presence or absence of *UBP8* and *UBP10* (Fig. 4A and S4A). Protein levels of Rad6, Bre1, and Rtf1 in all strains were similar to WT (Fig. S4B).

To test if the R6Q and R6A, M10A substitutions disrupt the response of Rad6 to the HMD, we purified Rad6-R6A, M10A and Rad6-R6Q proteins (Fig. S4C) and tested their activity in the reconstituted H2B ubiquitylation assay. Both mutant proteins were severely defective in responding to the HMD yet retained significant basal activity albeit at a lower level than WT (Fig. 4B-C, compare hatched to solid bars; and Fig. S4D). In agreement with the severe loss of global H2BK123ub (Fig. 4A), the *rad6-R6Q* and *rad6-R6A, M10A* mutants exhibit a telomeric silencing defect (Fig. 4D). However, when examined for UV sensitivity (Fig. 4E), N-end rule defects (Fig. 4F), and tolerance to oxidative stress (Fig. S4E), both mutants appear nearly indistinguishable from WT. Finally, given the reported links between H2BK123ub and DNA repair (49), we wanted to determine the level of UV sensitivity that could be ascribed to loss of H2BK123ub. Accordingly, we tested the UV sensitivity of strains containing genomic mutations in different H2B K123 ubiquitylation factors (Fig. S4F). This group included *rad18D* and *rad6D* strains as a reference for high sensitivity to UV (50). In addition to *rad6-R6Q* and *rad6-R6A, M10A* mutants, we tested a strain deleted for *BRE1, a* strain expressing H2B-K123R from both H2B-encoding genes (*htb1-K123R htb2-K123R*), and an *rtf1-108-110AAA* strain, which contains a triple amino acid substitution within the HMD (36) at the position of crosslinking with Rad6 (33). Compared to *rad6Δ* and *rad18Δ*, none of the other H2BK123ub-defective mutants show comparable levels of UV sensitivity (Fig. S4F). Collectively, our results highlight the key contributions of specific amino acids within the Rad6 N-terminal helix to a functional Rad6-HMD interface and H2BK123 ubiquitylation.

### Extensive overlap between the transcriptomes of *rad6* and HMD point mutants

To more comprehensively assess the extent to which the Rad6-R6A, M10A and R6Q mutants are specific to the H2BK123ub pathway, we utilized RNA sequencing (RNA-seq) as a sensitive phenotyping method. RNA-seq was performed on rRNA-depleted steady-state RNA from WT control strains and a set of integrated mutants: *rad6Δ*, H2B-K123R, *rtf1-108-110AAA, rad6-R6Q* and *rad6-R6A, M10A. Kluyveromyces lactis* was used as a spike-in control for data normalization (see Methods). We hypothesized that if the proposed Rad6-HMD interaction interface is specific to the H2BK123ub pathway, then mutating residues on either side of the interface will yield highly similar effects on the transcriptome, which will also phenocopy a H2B-K123R mutant.

All mutants except the H2B-K123R strain bear a 13xMyc C-terminal tag on *RAD6* and a 3xHA N-terminal tag on *RTF1*. Therefore, RNA-seq was performed on two WT strains (*RAD6 RTF1* and *RAD6-13xMyc 3x-HA-RTF1*) and two *rad6Δ* strains (*rad6Δ RTF1* and *rad6Δ 3xHA-RTF1*) to provide matched WT references for appropriate experimental samples and identify any tag effects. For all strains, results obtained from three biological replicates were highly reproducible (Fig. S5A). Differential expression analysis using DEseq2 (51) was performed on transcripts separately mapped to either the sense or antisense strands of coding regions. Principal Component Analysis (PCA) of the RNA-seq results showed WT strains clustering the furthest from *rad6Δ* strains with reference to PC1, while other H2BK123ub-defective mutants grouped between the two extremes (Fig. S5B). Comparisons between the two WT strains or the two *rad6*Δ strains revealed that less than 1% of all transcripts were differentially regulated due to the tags on *RAD6* and/or *RTF1* (p-adjusted ≤0.05) (Fig. S5C). Nevertheless, the comparison of WT strains revealed slight background differences within a narrow fold-change range and above the p-adjusted value 0.05 (Fig. S5C). Overall, these results justify further direct comparisons between the mutants.

Relative to their respective WT, all mutants displayed a large number of differentially expressed sense and antisense transcripts (Fig. 5A-B, top rows). The observed increase in antisense transcripts among the H2BK123ub-defective mutants is in agreement with reports of increased antisense transcription in *S. pombe* H2B ubiquitylation mutants (8, 52), although to our knowledge this is the first time this effect has been reported for an *S. cerevisiae* integrated H2B-K123R double mutant. To assess the extent of overlap between their transcriptomes, we next performed direct mutant-to-mutant comparisons (Fig. 5A-B, bottom rows). Comparisons of the *rad6-R6A, M10A* and *rad6-R6Q* mutants to each other or to the *rtf1-108-110AAA* mutant revealed almost no differences in either the sense or antisense direction (Fig. 5A-B, bottom rows and Fig. S5D). In contrast, direct comparisons of the *rad6-R6A, M10A* or *rad6-R6Q* transcriptomes to the *rad6Δ* transcriptome revealed hundreds of differentially expressed sense transcripts that passed the statistical cutoff, with 16% of these falling outside a log2 fold-change range of −1 to +1 (Fig. 5A, bottom and Fig. S5D, top). Collectively, these mutant comparisons strongly support the high specificity of the *rad6-R6A, M10A* and *rad6-R6Q* point mutations for the Rad6-HMD interaction.

**Figure 5:**
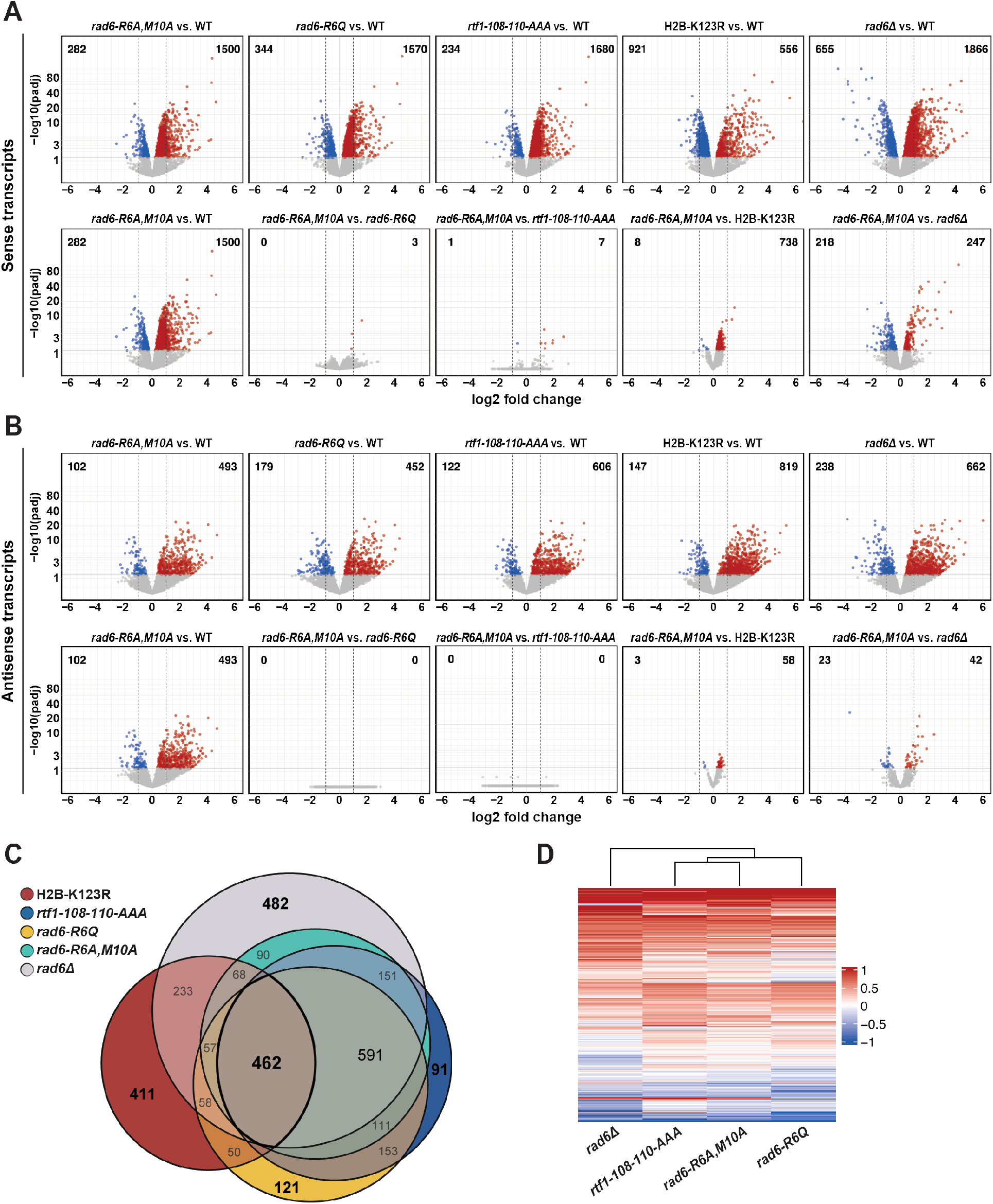
*rad6-R6Q* and *rad6-R6A, M10A* mutants phenocopy H2BK123ub-deficient mutants at the transcriptome level. (A) Top: Volcano plots displaying the differential expression analysis of sense transcripts in each mutant with respect to its corresponding WT control. All mutants were compared to the *3xHA-RTF1 RAD6-13xMyc* WT (*i.e.*, tagged WT) except H2B-K123R, which was compared to the *RAD6 RTF1* WT (*i.e.*, untagged WT). The *rad6*Δ mutant shown far right corresponds to the *rad6Δ 3xHA-RTF1* strain. The numbers of genes that pass a 0.05 p-adjusted value threshold whether up- (red) or down-regulated (blue) are shown at the top corners of each volcano plot. Bottom: Volcano plots of mutant-to-mutant comparisons. *rad6-R6A, M10A* was compared to the tagged WT and all the other mutants to emphasize the differences among the strains. Dashed vertical lines indicate the −1 to +1 range of log2 fold changes. Horizontal cutoff line represents the 0.05 p-adjusted (padj) value. (B) Same analysis as in A but based on the antisense transcripts only. (C) Venn diagram showing the overlap in the sets of differentially regulated sense transcripts (up and down) in each mutant relative to its corresponding WT. A significance threshold of p-adjusted value ≤ 0.05 was used. (D) Heatmap showing hierarchical clustering of the indicated mutant strains based on the log2-fold change values of the sense transcripts in the mutants with reference to WT.

When viewed as a Venn diagram, extensive overlap is also apparent among the differentially regulated sense transcripts in the *rad6* and *rtf1* point mutants relative to their WT controls (Fig. 5C). Consistent with the direct mutant-to-mutant comparisons, these genes represent only a subset of the differentially expressed genes observed in the *rad6Δ* strain. As expected, given the importance of the HMD-Rad6 interaction for H2B ubiquitylation, the transcriptome of the H2B-K123R mutant largely overlapped with the transcriptomes of the other mutants. However, this analysis also revealed differentially expressed transcripts unique to the H2B-K123R mutant. Although these transcriptional differences might reflect ubiquitylation-independent functions of K123, they appear to be dominated by small effects and more than 94% of them in the mutant-to-mutant comparisons are within a log2 fold-change range of −1 to +1 (Fig. 5A-B, bottom; Fig. S5D). Finally, hierarchical clustering based on differentially expressed sense transcripts highlights the close relationship among all three interface mutants and their distinction from the *rad6Δ* mutant (Fig. 5D). Overall, our RNA-seq results circumvent the limitations posed by single-phenotype tests and strengthen our conclusion that specific substitutions in the Rad6 N-terminal helix uniquely impair the functional interaction between Rad6 and the HMD.

## Discussion

In this study, we showed that the N-terminal helix of Rad6 is critical for its interaction with the Rtf1 subunit of Paf1C and for ubiquitylation of H2BK123, a conserved modification that broadly impacts chromatin structure, transcription, and a cascade of histone modifications. Our previous site-specific crosslinking experiments with BPA-substituted Rtf1 revealed a direct interaction between Rad6 and the HMD *in vivo* (33). Here, through site-specific crosslinking, genetic analysis, and *in vitro* H2B ubiquitylation assays, we identified Rad6 residues R6 and M10 as being especially important for and specific to the Rad6-HMD interaction. In support of this conclusion, mutation of either side of the proposed Rad6-HMD interface yielded strikingly similar transcriptomic profiles that were distinct from those of a *rad6Δ* strain. Our findings provide new mechanistic insights underlying Paf1C-stimulated co-transcriptional ubiquitylation of H2B.

The N-terminal helix of Rad6 is highly conserved and has been implicated in multiple Rad6 functions. Although it does not eliminate nuclear localization or charging by the E1 *in vitro*,a mutation that deletes the N-terminus of Rad6 (*rad6Δ1-9*) causes defects in the N-end rule pathway and interaction with Ubr1, H2BK123ub and downstream H3 methylation, telomeric silencing, Ty1 element transposition, and UV tolerance (9, 22, 50, 53). Our mutational studies identified specific amino acid substitutions in the Rad6 N-terminal helix that cause a range of H2BK123ub and telomeric silencing defects yet have little effect on UV sensitivity, N-end rule protein degradation or response to oxidative stress. One possible explanation for our results considering the pleiotropic effects of the *rad6Δ1-9* mutant is that the HMD-Rad6 interaction, and hence H2BK123ub, is particularly sensitive to the integrity of the Rad6 N-terminal helix. Other proteins that require an intact Rad6 N-terminal helix for their functions might interact with other regions of Rad6 and, therefore, are relatively resistant to the effects of single amino acid substitutions in the N-terminal helix. Notable among the substitutions we identified in our screen are Rad6-R6A, M10A and Rad6-R6Q, a cancer-associated change, which severely reduce deposition of H2BK123ub *in vivo* and stimulation by the HMD *in vitro*. Consistent with our results, a previous genetic screen identified a double mutant (*rad6-P4L, A5V*) that was partially defective in telomeric silencing but not N-end rule function or UV tolerance (53), although the mechanistic basis for the loss of telomeric silencing was undefined. The requirement of the N-terminal helix for various Rad6 functions, together with our discovery of N-terminal mutants that are specifically defective in H2BK123ub and stimulation by the HMD, suggests that this helix serves as a hub for multiple Rad6 binding partners.

Reported interactions involving the Rad6 N-terminal helix include interactions with Uba1 (E1) and Ubr1, the E3 for N-end rule proteolysis (43, 50, 54, 55). Rad6 R7, R8 and R11 constitute an E1 binding motif that is conserved among E2 proteins (55, 56). In the Rad6 structure, R8 is engaged in intramolecular hydrogen bonding that stabilizes the structural fold (44). Our *in vitro* data showing near complete loss of basal H2B ubiquitylation activity for the Rad6-R8A and Rad6-R11A mutant proteins are consistent with the involvement of these residues in protein stability and/or E1 binding. However, because Rad6_*Δ*1-9_ can be charged by the E1 *in vitro* (50), residues distal to the Rad6 N-terminal region must also contribute to the Rad6-E1 interaction (55). Unlike *rad6-R8A*, which is hypomorphic for many of the phenotypes tested (Fig. S2D), *rad6-R11A* appears to be selectively defective in H2BK123ub, suggesting this residue functionally or physically interacts with a key member of the H2B ubiquitylation machinery, possibly Bre1 (Fig. S3B). Recent cryo-EM structures show that Rad6 R6 is part of an extensive interaction interface (823.3 Å^2^) with Ubr1 (54). In the N-end rule assay used here, the *rad6-R6A, rad6-R6Q* and *rad6-R6A, M10A* mutants behave similar to WT, arguing that substitution of R6 alone is insufficient to severely compromise the interaction with Ubr1. In the context of existing literature, our data suggest that residues R6 and M10 are unlikely to confer strong effects on Rad6 binding to the E1 or Ubr1, although they might contribute to such interactions.

The striking similarities among the HMD (*rtf1-108-110AAA*) and Rad6 interface mutants (*rad6-R6Q* and *rad6-R6A, M10A*) on the transcriptome level, together with the phenotypic and transcriptomic distinction of these mutants from the *rad6*Δ strain, are indicative of the specificity of this interface for H2BK123ub. Consistent with this conclusion, the transcriptomic profile of the H2B-K123R mutant largely overlapped with those of the other mutants. We also observed an independent subset of transcripts specific to the H2B-K123R mutant, which might reflect loss of ubiquitylation-independent functions of K123 such as its role as a site for histone acetylation (57, 58). However, we cannot rule out that some of the deviation between the transcriptomic profiles among the mutants might stem from slight variation imposed by tagged alleles and/or spike-in normalization (Fig. S5A-C).

Although other contact points between Rad6 and Rtf1 might contribute, we believe that the Rad6-HMD interface proposed here is critical in targeting Rad6 to its histone substrate during transcription elongation. This likely happens in coordination with other Rad6-Paf1C contacts, including recently reported biochemical interactions of Rad6 with Ctr9 and Cdc73 (37). Additionally, Cdc73 and Rtf1, in an HMD-dependent manner, can accelerate ubiquitin discharge from Rad6 independently and in coordination with Bre1 *in vitro* (37). In light of other reported interactions between Bre1 and Rad6 (59, 60), Bre1 and the nucleosome (60), Bre1 and Paf1C (31), and the HMD and the nucleosome (61), it is evident that co-transcriptional H2B ubiquitylation is the product of a complex process entailing multivalent interactions. Future work is required to address the mechanistic and structural aspects that pose a critical requirement for the proposed Rad6-HMD interface and to pinpoint the order of events leading to efficient co-transcriptional H2B ubiquitylation *in vivo*.

## Materials and Methods

Yeast strains used in this study are derived from S288C (62) and listed in Table S2, and all plasmids are described in Table S3. Yeast cell extracts were made using a slightly modified NaOH extraction method (63), except for extracts in Fig. 3D that were made by TCA extraction as previously described (33). Proteins were resolved using SDS-polyacrylamide gel electrophoresis and analyzed by immunoblotting. *In vivo* BPA crosslinking and *in vitro* H2B ubiquitylation reactions were done essentially as described (33). Additional information and description of the statistical analysis and reproducibility is described in the Supporting Information (*SI Materials and Methods*).

### *In vitro* BPA crosslinking

For *in vitro* crosslinking followed by western analysis, BPA crosslinking of purified proteins was performed in conditions identical to the *in vitro* ubiquitylation reactions, except in Fig. 1B lane 1 and Fig. S1C lane 1 where buffer lacked ATP and contained 0.02 units of apyrase (Sigma A6535). Reaction mixtures were incubated for 10 min at 30°C before performing the crosslinking at 4°C for 30 min using long-wave UV light (365nm). A UVGL-55 handheld UV lamp was placed ~3 cm on top of the reaction mixture, which was dropped onto parafilm. The reaction mixture was then boiled in SDS loading buffer and run on a 12% SDS-polyacrylamide gel. For liquid chromatography mass spectrometry (LC-MS/MS) analysis, reactions contained 37 μg HMD_74-184-F108BPA_ and 16.25 μg V5-Rad6 in a volume of 105 μl. Buffer conditions were 8% glycerol, 7.2 mM Tris-Cl pH 8.0, 26 mM NaCl, and 0.33 mM β-mercaptoethanol. After 10 min incubation at 30 °C, the reaction volume was equally divided into 10 wells of a 96-well plate (Life Technologies #4346907) and covered with an optically clear adhesive seal (Thermo Fisher Scientific #AB-1170). The UV lamp was rested directly onto the plate and crosslinking was performed at 4°C for 30 min at 365 nm. The full reaction was then re-pooled, boiled in SDS loading buffer and run on a 12% SDS-polyacrylamide gel (Bio-Rad) followed by Coomassie staining. Gel slices were excised and processed for LC-MS/MS. MS methods have been previously described (64).

### RNA-seq and data analysis

*S. cerevisiae* and *K. lactis* (KL01) cells were grown in YPD, harvested at log phase (OD_600_ = 0.8–1.0; measured on an Eppendorf BioPhotometer) and mixed in a 9:1 ratio, respectively, according to the OD_600_ units. Total RNA was extracted from the spiked-in cell mixture using hot acid phenol (65). The quality of extracted RNA was assessed by gel electrophoresis on 1% TBE gels, and concentrations were determined by Qubit RNA HS kit (Invitrogen). DNase treatment of the RNA samples was done on-column using RNase-Free DNase Set (QIAGEN) followed by rRNA-depletion using *S. cerevisiae* riboPOOLs probes (siTOOLs Biotech). Sequencing libraries were prepared using NEBNext^®^ Ultra™ II Directional RNA Library Prep Kit for Illumina (NEB #E7760) coupled with AMPure XP bead purification (Beckman Coulter) with 18-22 ng of rRNA-depleted RNA as an input. Library quality and fragment distribution were assessed on an Agilent fragment analyzer. Samples were sequenced at the Health Sciences Sequencing Core at UPMC Children’s Hospital on a NextSeq 500 with 2 × 43-bp paired-end reads. Using STAR (v2.7.5a) aligner (66), all reads were first aligned to the *K. lactis* genome (Ensembl ASM251v1). Reads unmapped to *K. lactis* were designated for alignment to the *S. cerevisiae* S288C genome (Ensembl R64-1-1). Transcript counts were extracted from the BAM files using featureCounts (67). To perform the spike-in correction of *S. cerevisiae* counts, *K. lactis* read counts (mapping to sense strands only) were fed into DESeq2 (51) to estimate replicate-specific size factors. Estimated size factors were then applied to *S. cerevisiae* counts that correspond to proteincoding genes only. Genes with zero counts in all samples were filtered out before the differential expression analysis. To increase the sensitivity of detecting genotype-dependent significant differences, batch effects were mitigated by applying a multi-factor design in DESeq2 that includes the genotype and the batch of each sample (i.e., biological replicate number) consistent with how samples were processed in groups of the same biological replicate. PCA plots were derived from counts that underwent Variance Stabilizing Transformation (VST) (51). Tidyverse, Psych, ComplexHeatmap and eulerr R packages were used to produce the correlation plots, heatmap and Venn diagrams. All genomic data has been deposited at GEO under accession number GSE218023.

## Acknowledgments

We dedicate this manuscript to the memory of Ivan Belashov, who contributed to the early stages of this project during his research rotation in the Arndt lab. We are grateful to Jaehoon Kim for purified recombinant hE1 and yBre1 and to Song Tan for *X. laevis* mononucleosomes. We thank Sarah Hainer, Craig Kaplan, Miler Lee, Andrew Vandemark, Andrea Berman, members of their groups, and all members of the Arndt lab, especially Alex Francette, for many helpful suggestions. We thank Fred Winston for critical reading of the manuscript. This project used the University of Pittsburgh Health Sciences Sequencing Core at UPMC Children’s Hospital of Pittsburgh for genomic sequencing experiments and was supported in part by the University of Pittsburgh Center for Research Computing through the resources provided. The mass spectrometry work was supported, in part, by the Indiana Clinical and Translational Science Institute, funded in part by Award Number UL1TR002529 from the NIH, National Center for Advancing Translational Sciences, Clinical and Translational Sciences Award. Research conducted in the Arndt lab was supported by the National Institutes of Health (R01GM52593 and R35GM141964 to K.M.A).

