## supplemental materials for "Paf1 complex subunit Rtf1 stimulates H2B ubiquitylation by interacting with the highly conserved N-terminal helix of Rad6"

#### SI Materials and Methods

##### Yeast strains and genetic methods

Yeast strains used in this study are derived from S288C (1) and listed in Table S2. Strains were generated using standard techniques for genetic crosses and gene replacements (2). All PCR amplified regions for genomic integration or plasmid construction were confirmed by DNA sequencing. Transformations of yeast were performed using the lithium acetate-PEG method (3). Yeast cells were grown at 30°C in YPD or synthetic complete (SC) media with omission of appropriate nutrients to maintain plasmid selection (2). 2% glucose or galactose was used as the carbon source for appropriate experiments, as indicated.

The construction of the strain in which both copies of H2B contained a K123R substitution (i.e., integrated double mutant) was done as follows. First, a strain expressing *htb2-K123R* was created by *delitto perfetto* (4) in which the open reading frames for *HTA2-HTB2* were replaced with pCORE using homologous recombination in KY3694 and selecting for $Ura^+$, $G418^R$ colonies to create KY3733. Next, overlapping PCR was used to create a DNA fragment containing *HTA2-htb2-K123R* that replaced the pCORE fragment by selecting for $5FOA^R$ colonies and screening for $G418^S$. After sequencing the entire replacement region in the resulting strain, KY3761, a genetic cross was performed with KY2876, which contained *htb1-K123R* substitution to generate KY3781. Substitutions in *HTB2* were inferred to be in strains lacking *TRP1* and the *htb1-K123R* alleles were identified by restriction enzyme digestion as previously described (5). RNA-seq data confirmed the substitutions.

##### Plasmid construction

Plasmids were constructed using both traditional cloning techniques and Gibson assembly (New England Biolabs #E2611S). Plasmid descriptions are provided in Table S3. Point mutants were generated via PCR with primers containing the desired mutations, followed by Gibson assembly into the appropriate plasmid backbone. pQLinkN-HMD$_{74\text{-}184\text{-}F108BPA}$ (pKB1441) was constructed by amplifying the HMD$_{74\text{-}184\text{-}F108BPA}$ sequence from a derivative of pKB1228, in which an amber codon is in place of codon 108 of *RTF1*. A *BamH*I restriction site and a C-terminal 7xHis tag were added to the sequence by PCR and the resulting fragment was cloned into pQLinkN (6) using the *BamH*I site.

**Serial dilution spot assays**

Yeast transformants were grown to saturation in selective media and then pelleted and resuspended to $OD_{600}$=1.0 in sterile dd$H_2O$. For telomeric silencing assays, five-fold serial dilutions of KY3391 transformants were spotted onto medium containing 5FOA (Goldbio #F-230) or onto growth control medium and incubated at 30°C. For UV sensitivity assays, five-fold serial dilutions of KY2045 transformants were plated onto SC-Trp plates, then exposed to the indicated UV dosage using a UV crosslinker (Fisher Scientific FB-UVXL-1000) and incubated in the dark at 30°C. The same was done to test the UV sensitivity of the integrated mutants in Fig. S4F, except that they were plated onto SC (complete) plates and then exposed to different UV dosages as described. To test response to oxidative stress, KY2045 transformants were grown to log phase ($OD_{600}$=0.3-0.6). Cells were then pelleted and resuspended in fresh medium containing 2.5 mM $H_2O_2$. After incubation for 45 min at 30 °C on a rotating drum, all cultures were normalized back to an $OD_{600}$ of 0.2, and five-fold serial dilutions were plated. Time of imaging relative to plating is noted in the figure legends.

**Preparation of yeast cell extracts**

Yeast cells were grown to mid-log phase ($OD_{600}$=0.7-1.3) and 10 or 25 $OD_{600}$ units of cells were pelleted and flash frozen in liquid nitrogen before processing. Extracts were made using a slightly modified NaOH extraction method (7). Briefly, cell pellets, corresponding to 10 $OD_{600}$

units, were thawed and resuspended in 400 µl water; then 400 µl 0.2M NaOH was added. Cells were incubated for 5 min at room temperature and centrifuged at 16,000xg for 5 min. Supernatant was discarded, and pellets were resuspended in 200 µl SDS loading buffer. Samples were boiled for 5 min and clarified by spinning at 16,000xg for 5 min at room temperature before flash freezing in liquid nitrogen. Extracts for Fig. 3D were made by a TCA extraction method (8).

**Western blotting**

SDS-polyacrylamide gels were transferred to nitrocellulose, then rinsed in water and Ponceau stained to verify efficient transfer. Blots were destained in TBST, then blocked for 1 hr in 5% milk in TBST. Primary antibodies were α-V5 (Invitrogen #R960-25; 1:1500 dilution), α-Rtf1 (1:2500) (9), α-H2B K120ub (Cell Signaling #5546; 1:1000), α-H2B (Active Motif #39237; 1:3000), α-G6PDH (Sigma #A9521; 1:20,000), α-Myc (generous gift of John Woolford; 1:100), α-H3K4me3 (Active Motif #39159; 1:2000), α-H3K79me2/3 (Abcam #ab2621; 1:1000), α-H3 (1:15,000) (5), α-H2A (Active Motif #39235; 1:5000), and α-HA (Santa Cruz #sc-7392; 1:3000). With the exception of α-H3 (1% milk in TBST), primary incubations were performed in 5% milk in TBST for 2-3 hr at room temperature or overnight at 4°C. Following primary incubations, blots were rinsed in TBST and secondary incubations were performed for 1-2 hr using 1:5000 dilutions (in TBST) of the appropriate HRP-conjugated secondary antibodies (either donkey anti-rabbit or sheep anti-mouse; GE Healthcare). Some dilutions of secondary antibodies contained 2-5% milk to increase specificity, particularly when primary incubations were performed overnight. Visualization was achieved using SuperSignal™ West PicoPLUS chemiluminescent substrate (Thermo #34579) and a Bio-Rad ChemiDoc XRS+ imager. Image analysis and quantifications was performed using BioRad Image Lab 6.0.1 software.

***In vivo* BPA (p-benzoyl-L-phenylalanine) crosslinking**

Strain KY2798 (*3xHA-RTF1 rad6Δ::URA3*) was co-transformed with pLH157/LEU2 (8), which encodes the tRNA/tRNA synthetase needed for BPA incorporation, and a 2µ plasmid expressing C-terminally 13xMyc-tagged Rad6-P4BPA (pKB1585), Rad6-R7BPA (pKB1586) or Rad6-F13BPA (pKB1514) upon amber codon suppression (10). Cells were grown to early log phase in SC-Leu-Trp medium supplemented with 1 mM BPA (Bachem, #F-2800) in 1M HCl, which was neutralized with an equal volume of 1M NaOH directly after addition to the medium. Cells (10 $OD_{600}$ units) were pelleted and resuspended in 1 ml of sterile dd$H_2O$. The cell suspension was placed in a sterile petri dish allowing the surface tension to form a circular drop. A UVGL-55 handheld UV lamp (UVP) was positioned 2 cm from the cells, which were exposed to UV-induced crosslinking at a wavelength of 365 nm for 10 min. For western analysis, collected cells were used to prepare TCA extracts as previously described (8).

***In silico* structural modeling**

Modeling of the HMD-Rad6 interaction was performed using the ClusPro 2.0 webserver (https://cluspro.bu.edu/) (11–15). Structures of Rad6 (PDB: 1AYZ, chain A) and $HMD_{74–139}$ (PDB: 5E8B, chain A) (8, 16) were used as receptor and ligand. Based on the *in vitro* BPA crosslinking and LC-MS/MS results, a distance restraint of a maximum of 10 Å was applied between HMD F108 and each of Rad6 L9, M10, R11 and D12, with 100% required percentage of restraints. The model shown in Fig. 3C satisfied the following criteria: 1) Rtf1 108 and Rad6 M10 are in close proximity, 2) Rad6 R6 is proximal to the HMD to be consistent with the biochemical data, and 3) other Rtf1 residues that fail to support *in vivo* BPA crosslinking (E83, K85, R92 and E96) (8) are within a non-crosslinking distance to the nearest Rad6 residues (>10 Å). In addition, full-length protein sequence of Rad6 and the sequence corresponding to Rtf1 $HMD_{73-139}$ was used for complex prediction using Alphafold2-multimer through ColabFold (17–22). PyMOL (23) was used to generate representative views of the top-ranking model from each method.

**N-degron reporter assay**

KY3392 (*rad6*Δ) or KY3389 (*rad6*Δ *ubr1*Δ) were co-transformed with the indicated *RAD6* plasmids and a derivative of pUB23 (pKB1526) (24). The latter plasmid encodes a ubiquitin-Arg-β-galactosidase fusion protein from which the ubiquitin is cleaved *in vivo* and the N-terminal Arg is exposed. Cells were grown to mid-log phase in SC-Trp-Ura medium containing 2% galactose. Extracts were made by bead beating in breaking buffer (100 mM Tris-Cl pH 8.0, 20% glycerol, 1mM DTT, Halt$^{TM}$ Protease inhibitors (Thermo Fisher Scientific #78430) followed by flash freezing in liquid nitrogen. Extracts concentrations were determined by Bradford assay (Bio-Rad). The volume corresponding to 15 µg protein per sample was adjusted to 20 µl with breaking buffer, and then 80 µl of Z-buffer (100 mM $Na_2HPO_4$, 40 mM $NaH_2PO_4$, 10 mM KCl, 1 mM $MgSO_4$, 0.27% BME) was added. Reactions were started by adding 20 µl ONPG (4 mg/ml) per well of a 96-well plate (Thermo Fisher Scientific #655180), and $OD_{412}$ readings were collected once per minute for 2 hr using a BioTek Cytation5 plate reader. $V_{max}$ for each sample was determined using Gen5 software. Unless stated otherwise, the $V_{max}$ value of each biological replicate was the average of two measurements (technical duplicate). To determine the activity relative to WT, an average $V_{max}$ of all readings from WT *RAD6* extracts was used as the baseline reference, and the $V_{max}$ values corresponding to the mutants were divided by this baseline reference.

**Protein purification**

Rtf1 $HMD_{74-184}$ protein and His-pK-HA-Ubiquitin were purified as described (8). To purify 6xHis-MBP-TEV-V5-Rad6, 8xHis-TEV-V5-Rad6 and 8xHis-TEV-Rad6 or the point mutant derivatives of the latter, *E. coli* BL21 (DE3) Codon+ (RIPL) cells were transformed with the appropriate expression plasmids (Table S3) and grown in LB medium supplemented with 100 µg/ml ampicillin and 34 µg/ml chloramphenicol. Rad6 expression was induced by addition of IPTG

(200 µM final concentration) to cells in log phase. Cells were grown shaking (160 rpm) at room temperature overnight. After induction, purification was done by nickel affinity chromatography (QIAGEN Ni-NTA agarose #30210) followed by an overnight digestion with His-tagged TEV protease. The TEV protease and uncleaved proteins were removed by a second nickel affinity chromatography step followed by ion exchange chromatography using HiTrap-SP HP and HiTrap-Q FF columns (GE Healthcare). The proteins were then dialyzed overnight into storage buffer (10% glycerol, 25 mM Tris pH 8.0, 150 mM NaCl, 1 mM β-mercaptoethanol) and concentrated using a 5 kDa molecular-weight cutoff Vivaspin concentrator (Sartorius VS15T11). Flag-hE1 and Flag-yBre1 were generously provided by Jaehoon Kim, and recombinant *X. laevis* nucleosomes were generously provided by Song Tan. For $HMD_{74\text{-}184\text{-}F108BPA}$–7xHis protein purification, BL21(DE3) cells were co-transformed with pKB1441 (pQLinkN-$HMD_{74\text{-}184\text{-}F108BPA}$) and pEVOL-pBpF (25) (sourced from Addgene) and grown in Terrific Broth supplemented with 100 µg/ml ampicillin, 34 µg/ml chloramphenicol, and 500 µM BPA. Induction of the tRNA synthetase (on pKB1444) was achieved by addition of arabinose to 0.2% (w/v) final concentration, and HMD expression was induced by addition of IPTG to 250 µM final concentration. During induction, cells were grown at 37 °C for 3-4 hrs. Following cell lysis, the protein was purified by nickel affinity chromatography followed by ion exchange chromatography. The protein was then dialyzed overnight into storage buffer (30% glycerol, 20 mM Tris pH 8.0, 50 mM NaCl, 1 mM β-mercaptoethanol) and concentrated using a 5 kDa MWCO concentrator.

***In vitro* H2B ubiquitylation reactions**

Reactions were performed essentially as described in (8). Briefly, 10 µl reactions contained 50 ng Flag-hE1, 1.4 µg His-pK-HA-Ubiquitin, and 2.5 µg *X. laevis* nucleosomes in buffer containing 50 mM Tris-Cl pH 7.9, 5 mM $MgCl_2$, 2 mM NaF, 0.4 mM DTT, and 4 mM ATP. Reactions contained 100 ng Flag-yBre1, 100 ng Rad6, and 400 ng (Fig. 3A-B) or 200 ng (Fig. 4B-C) of

$HMD_{74-184}$. Reactions were incubated for 30 min at 30°C before boiling in SDS loading buffer. Reactions were run on 15% SDS-polyacrylamide gels and assessed by western blotting. For quantification, the intensity of the signals for both H2BK123ub and H2A was quantified using Image Lab Software (version 6.1). H2BK123ub signal was normalized to the H2A signal from the same lane. Normalized H2BK123ub signal corresponding to the reaction containing WT Rad6 with Bre1 and lacking $HMD_{74-184}$ (WT middle lane) was used as a baseline reference. Accordingly, H2BK123ub fold stimulation relative to WT was estimated by dividing the normalized H2BK123ub signal in all other lanes by this baseline reference.

**Mass Spectrometry Data Analysis**

Raw mass spectrometry files (mzXML) were processed using Crossfinder version 1.1 (26, 27). The following settings were used consistently for all samples. The crosslinker was set as BPA with a monoisotopic mass of 251.0946 Da on HMD at the site of BPA incorporation. Trypsin cleavage was used with a maximum missed cleavage of 2. Modifications for variable methionine oxidation (0 for unoxidized, 15.9949 for oxidized, or 31.9898 Da for doubly oxidized) and fixed cysteine oxidation (57.021464 Da) were included. The settings for allow.elim and allow.loop were set to “FALSE” while MS tolerances were left blank. The MS2 rank cutoff was set at 12 and the MS2 rank mass window was set at 50. Neutral water loss for both b and y ions was set at -18.0106 Da, and the minimal amino acid length was 4. The following filters were used for the data analysis: filers.nFrags_total = 6, filters.nXLfrags_total = 0, filters.nFrags_perPep = 2, filters.nUnqFrags_perPep = 0, filters.nConsecFrags_perPep = 0, filters.fracTIC = 0.05, filters.minScore_total = 200, and filters.score_rel = 90. All samples were searched against custom made FASTA databases with the proteins in question included (yeast Rad6 and Rtf1). A custom made FASTA database with common contaminants was also included.

**Statistical analysis and reproducibility**

All *in vivo* experiments were performed in a minimum of three replicates, including the RNA-seq experiment. For all experiments involving live cell cultures, each biological replicate is derived from a single colony of a particular strain. For experiments done with transformants, the three biological replicates represent independent transformants. Spot assays were done in at least two technical replicates for each of the three biological replicates. For *in vitro* experiments, technical replicates were done using the same sources of the recombinant proteins and were performed a minimum of two times, with the exception of the *in vitro* crosslinking-mass spectrometry experiment in which one representative replicate was used to guide subsequent mutagenesis. The p-values of western blot quantification were calculated based on unpaired two-tailed Student's t-test. Significance of the N-end rule measurements was assessed using unpaired one-way ANOVA followed by a Dunnett's multiple comparison test with reference to the WT. For RNA-seq differential expression analysis, Wald test was applied through DESeq2 to call significant fold changes with a threshold of adjusted p-value $\leq 0.05$. Graphs other than ones in Figures 5 and S5 were produced in Prism.

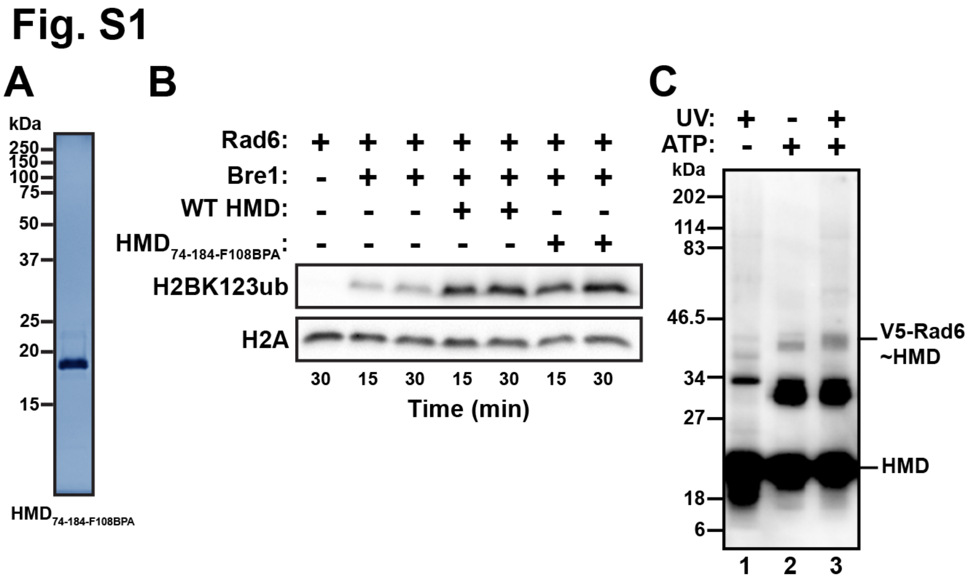


**Figure S1: HMD$_{74\text{-}184\text{-}F108BPA}$ protein used for *in vitro* crosslinking stimulates H2BK123ub *in vitro*.** (A) False-colored coomassie-stained gel displaying the purified HMD$_{74\text{-}184\text{-}F108BPA}$ protein used for *in vitro* crosslinking experiments. (B) Western blot analysis of the products from a reconstituted H2B ubiquitylation reaction. Purified HMD$_{74\text{-}184\text{-}F108BPA}$ stimulates H2BK123ub levels comparable to purified WT HMD$_{74\text{-}184}$. (C) Western blot showing the same reactions as in Fig. 1B but probing with Rtf1 antisera, which detects the HMD$_{74\text{-}184}$. The position of the V5-Rad6~HMD$_{74\text{-}184\text{-}F108BPA}$ crosslinked species is indicated and runs close to a low-intensity band representing an uncrosslinked species (compare lanes 2 and 3). Other reaction products are likely ubiquitylated forms of the HMD (lanes 2 and 3) or a crosslinked dimer of the HMD (lanes 1-3, ~34 kDa).

### Fig. S2

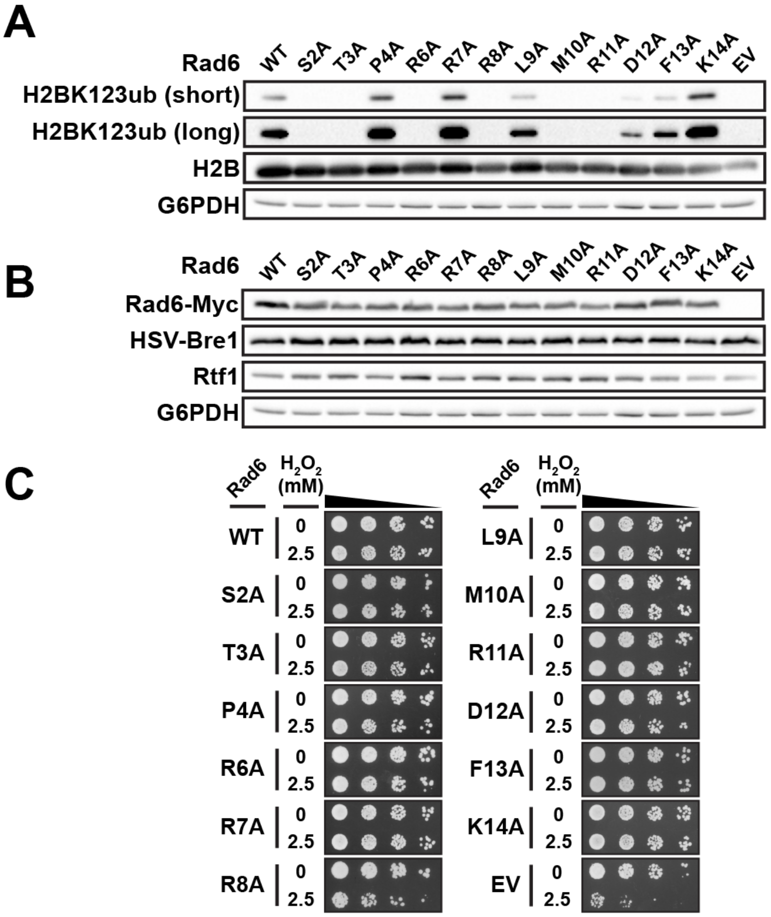


D

| | | Rad6 mutant | | | | | | | | | | | | |
|---|---|---|---|---|---|---|---|---|---|---|---|---|---|---|
| Tested phenotypes | WT | S2A | T3A | P4A | R6A | R7A | R8A | L9A | M10A | R11A | S2A | F13A | K14A | EV |
| *In vivo* bulk H2BK123ub | +++ | + | - | +++ | - | +++ | - | ++ | - | + | ++ | +++ | +++ | - |
| *In vivo* bulk H3K4me3 and H3K79me2/3 | +++ | ++ | ++ | +++ | + | +++ | + | +++ | + | ++ | +++ | +++ | +++ | - |
| Telomeric silencing | +++ | - | - | +++ | - | +++ | - | ++ | - | - | + | + | ++ | - |
| Sensitivity to UV | +++ | +++ | +++ | +++ | +++ | +++ | - | +++ | +++ | +++ | ++ | +++ | +++ | - |
| Protein degradation | +++ | +++ | +++ | +++ | +++ | +++ | ++ | +++ | +++ | +++ | +++ | +++ | +++ | - |
| Sensitivity to oxidative stress | +++ | +++ | +++ | +++ | +++ | +++ | ++ | +++ | +++ | +++ | +++ | +++ | +++ | - |

Phenotypic score: WT +++ ++ + - *rad6Δ*

**Figure S2: Analysis of protein levels and additional phenotypes for the *rad6* point mutants.** In A and B, a *rad6*Δ *1xHSV-BRE1* strain (KY3551) was transformed with WT or mutant *rad6* plasmids or empty vector (EV). (A) Western analysis showing that the 1xHSV tag on Bre1 does not affect the pattern of bulk H2BK123ub levels in the *rad6* alanine scanning mutants (compare to Fig. 2A). Short and long exposures of the same H2BK123ub blot are shown. (B) Western analysis showing bulk protein levels of Rad6-13xMyc, 1xHSV-Bre1 and Rtf1 in the *rad6* alanine scanning mutants. (C) Five-fold serial dilution spot assay to assess cell growth under oxidative stress. Transformants of a *rad6*Δ strain (KY2045) were treated with 2.5 mM $H_2O_2$ for 45 min at early log phase and plated onto SC-Trp medium without $H_2O_2$. The representative images show growth on day two relative to the time of plating. (D) Summary of *rad6* point mutant phenotypes compared to WT and *rad6*Δ (EV) control strains.

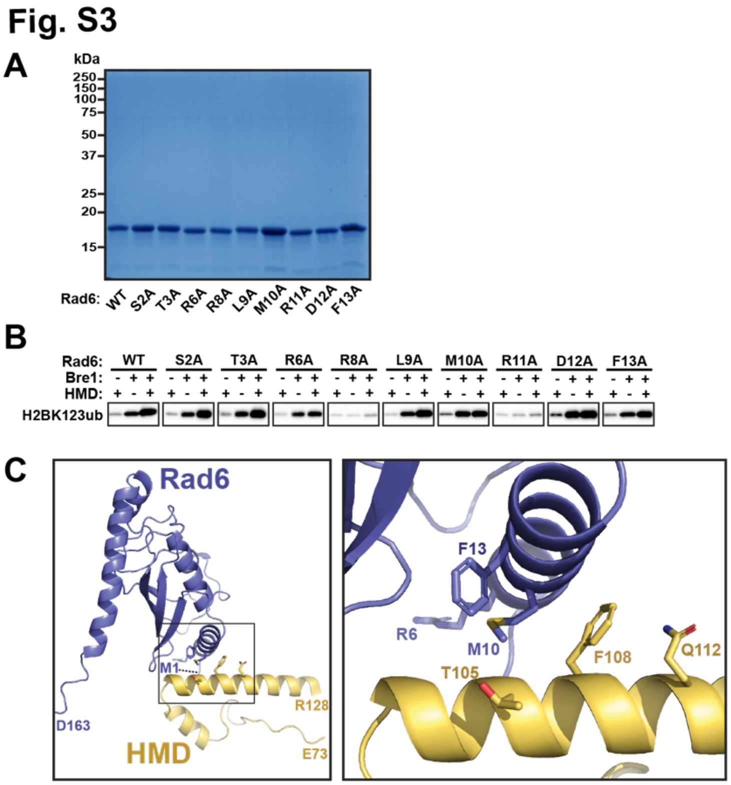


**Figure S3: Characterization of recombinant Rad6 mutant proteins** (A) False-colored coomassie-stained gel of purified Rad6 proteins. (B) Long exposures of western blots shown in Fig. 3A to emphasize H2BK123ub levels in low-signal lanes. (C) Alphafold2-multimer prediction of the Rad6-HMD complex. The inset in the left panel is expanded on the right.

#### Fig. S4

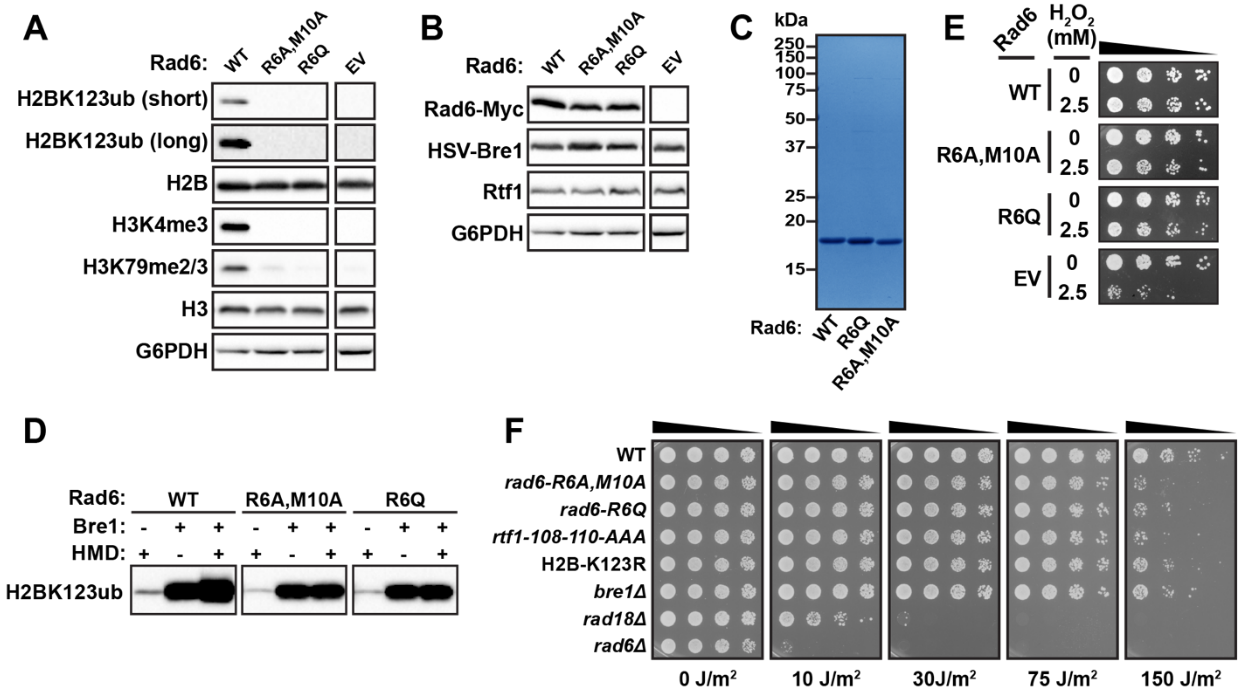


**Figure S4: Characterization of *rad6-R6A, M10A* and *rad6-R6Q* mutants.** (A) Western blot showing H2BK123ub and H3 methylation levels in transformants bearing the *1xHSV-BRE1* allele. The tag on Bre1 does not alter histone modification patterns in the *rad6* mutants (compare to Fig. 4A). Short and long exposures of the same H2BK123ub blot are shown. EV = empty vector. G6PDH serves as a loading control. (B) Western blot showing bulk protein levels of Rad6-13xMyc, 1xHSV-Bre1 and Rtf1 in *rad6-R6A, M10A* and *rad6-R6Q* mutants. For both panels A and B, bands in each row come from the same blot, including the EV lane; lanes were re-arranged for clarity. (C) False-colored coomassie-stained gel displaying Rad6-R6A, M10A and Rad6-R6Q purified proteins. (D) Long exposures of blots shown in Fig. 4B. (E) Five-fold serial dilution spot assay to assess cell growth under oxidative stress. Transformants of a *rad6*Δ strain (KY2045) were treated with 2.5 mM $H_2O_2$ for 45 min at early log phase and plated onto SC-Trp medium without $H_2O_2$. Representative images show growth on day two relative to the time of plating. (F) Five-fold serial dilution spot assay of WT (KY3751) and strains with the following mutations integrated to replace the endogenous gene: *rad6-R6A, M10A* (KY3751), *rad6-R6A* (KY4018), *rtf1-108-110AAA* (KY3748), H2B-K123R (KY3781), *bre1Δ* (KY954), *rad18Δ* (KY2439) and *rad6Δ* (KY3737). Cells were grown to saturation in YPD liquid medium and then plated to SC-complete medium, followed by exposure to various doses of UV irradiation. Representative images show growth on day two relative to the time of plating.

#### Fig. S5

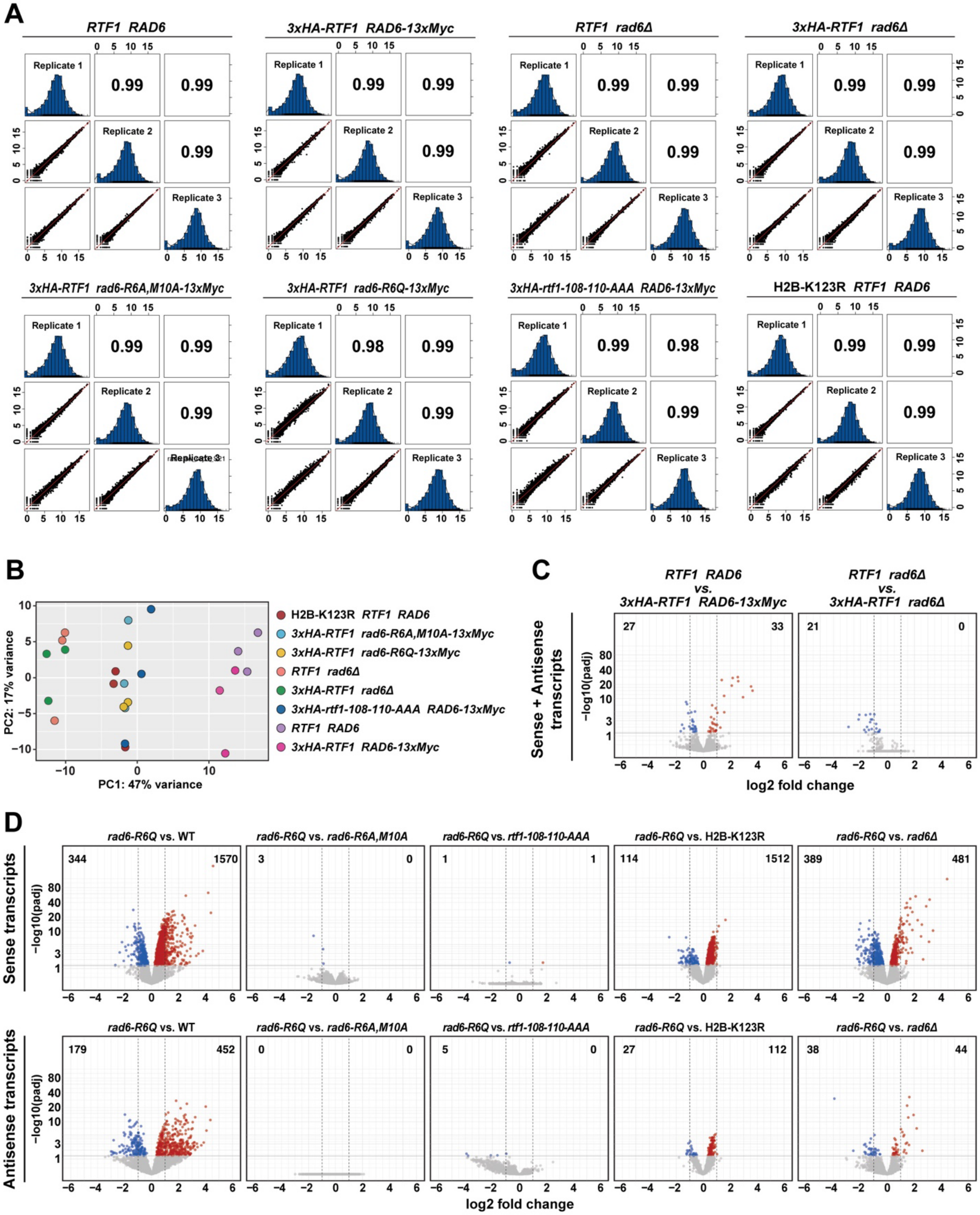


**Figure S5: Analysis of RNA-seq replicates and *rad6-R6Q* mutant differential expression.** (A) Correlation analysis of RNA-seq biological replicates indicate high reproducibility. For comprehensive assessment, counts from both sense and antisense transcripts were included.

Pearson's correlation coefficients are indicated inside the squares. Bivariate scatter plots (i.e., XY plots) showing the correlation between log2-transformed spike-in normalized counts between each pair of replicates as indicated. Frequency histograms show the distribution of the log2-transformed normalized counts within each replicate. (B) PCA plot of all biological replicates using normalized counts from both sense and antisense transcripts. (C) Volcano plots comparing tagged and untagged WT or *rad6*Δ strains as indicated. Dashed vertical lines indicate the -1 to +1 range of log2 fold changes. Horizontal cutoff line represents the 0.05 p-adjusted (padj) value. The numbers of genes that pass 0.05 p-adjusted value threshold whether up- (red) or down-regulated (blue) are shown at the top corners of each volcano plot. (D) Volcano plots of the sense or antisense differential expression analysis of *rad6-R6Q* compared to WT and all other mutants (similar to Fig. 5A-B, bottom rows).

**Table S1: Mass spectrometry results**

V5-Rad6 protein sequence*:
*GHMAS* **GKPIP NPLLGLDST** *G SAS* MSTPARR RLMRDFKRMK EDAPPGVSAS PLPDNVMVWN AMIIGPADTP YEDGTFRLLL EFDEEYPNKP PHVKFLSEMF HPNVYANGEI CLDILQNRWT PTYDVASILT SIQSLFNDPN PASPANVEAA TLFKDHKSQY VKRVKETVEK SWEDDMDDMD DDDDDDDDDD DDEAD

*Linker sequences: italics. V5 tag: bold. Rad6 sequence: underlined.

| Region | Residue | Peptide Spectrum Matches (PSMs) |
|---|---|---|
| V5 tag | A4 | 1 |
| | S5 | 1 |
| | G6 | 1 |
| | K7 | 1 |
| | P8 | 1 |
| | I9 | 2 |
| | P10 | 1 |
| | N11 | 1 |
| | P12 | 3 |
| | L13 | 3 |
| | L14 | 8 |
| | G15 | 9 |
| | L16 | 12 |
| | D17 | 15 |
| | S18 | 17 |
| | T19 | 19 |
| | G20 | 19 |
| | S21 | 19 |
| | A22 | 18 |
| | S23 | 16 |
| Rad6 N-terminal region | M1 | 11 |
| | S2 | 2 |
| | T3 | 1 |
| | P4 | 1 |
| | A5 | 1 |
| | R6 | 1 |
| | R7 | 1 |
| | R8 | 1 |
| | L9 | 19 |
| | M10 | 38 |
| | R11 | 28 |
| | D12 | 13 |
| | K14 | 1 |
| | R15 | 3 |
| Rad6 C-terminal region | K134 | 2 |
| | S135 | 1 |
| | Q136 | 1 |

| | | |
|---|---|---|
| | Y137 | 2 |
| | V138 | 3 |
| | K139 | 3 |
| | R140 | 1 |
| | V141 | 1 |
| | K142 | 2 |
| | E143 | 1 |
| | T144 | 1 |
| | V145 | 2 |
| | E146 | 2 |

**Table S2: Yeast strains**

| Strain | *MAT* | *Genotype* |
|---|---|---|
| KY669 | a | *his3Δ200 lys2-128δ leu2Δ1 ura3-52 trp1Δ63* |
| KY954 | a | *lys2-128δ leu2Δ1 ura3-52 trp1Δ63 bre1Δ::kamMX4* |
| KY2045 | α | *his3Δ200 leu2Δ1 trp1Δ63 rad6Δ::KanMX* |
| KY2439 | a | *his3Δ200 leu2Δ1 trp1Δ63 rad18Δ::HIS3* |
| KY2798 | a | *his3Δ200 lys2-128∂ leu2∆1 ura3-52 trp1∆63 3xHA-RTF1 rad6Δ::URA3* |
| KY2876 | α | *his3∆200 leu2∆1 ura3∆0 trp1∆63 HTA1-htb1K123R (hta2-ht2b∆)::TRP1 GAL1p-FLO8-HIS3::KanMX TELVR::URA3* |
| KY3389 | α | *his3∆200 lys2-128∂ leu2∆1 ura3-52 trp1∆63 rad6∆::KanMX ubr1∆::HIS3* |
| KY3391 | α | *his3∆200 leu2∆1 ura3-52 trp1∆63 TELVR::URA3 rad6∆::KanMX* |
| KY3392 | α | *his3∆200 lys2-128∂ leu2∆1 ura3-52 trp1∆63 rad6∆::KanMX* |
| KY3551 | α | *his3∆200 lys2-128∂ leu2∆1 ura3-52 trp1∆63 rad6Δ::URA3 1xHSV-BRE1* |
| KY3694 | a | *his3Δ200 lys2-128δ leu2Δ1 ura3-52 trp1Δ63 hta2-htb2Δ::TRP1* |
| KY3733 | a | *his3Δ200 lys2-128δ leu2Δ1 ura3-52 trp1Δ63 hta2Δ::pCORE-KanMX-URA3::htb2Δ* |
| KY3737 | a | *his3∆200 lys2-128∂ leu2∆1 ura3-52 trp1∆63 3xHA-RTF1 rad6∆::KANMX* |
| KY3748 | a | *his3Δ200 lys2-128δ leu2∆1 ura3-52 trp1Δ63 3xHA-rtf1-108-110AAA RAD6-13xMYC::KanMX* |
| KY3751 | a | *his3Δ200 lys2-128δ leu2∆1 ura3-52 trp1Δ63 3xHA-RTF1 RAD6-13xMYC::KanMX* |
| KY3761 | a | *his3Δ200 lys2-128δ leu2Δ1 ura3-52 trp1Δ63 HTA2-htb2-K123R* |
| KY3781 | a | *his3∆200 lys2-128δ leu2∆1 ura3-52 trp1Δ63 HTA1-htb1K123R HTA2-htb2-K123R* |
| KY4010 | a | *lys2-128∂ ura3-52 trp1Δ63 rad6Δ::URA3 ubp8Δ::kanMX4 ubp10Δ::KanMX* |
| KY4017 | a | *his3Δ200 lys2-128δ leu2Δ0 ura3-52 trp1Δ63 3xHA-RTF1 rad6-R6A,M10A-13xMyc::KanMX* |
| KY4018 | a | *his3Δ200 lys2-128δ leu2Δ0 ura3-52 trp1Δ63 3xHA-RTF1 rad6-R6Q-13xMyc::KanMX* |
| KY4019 | a | *his3Δ200 lys2-128δ leu2Δ0 ura3-52 trp1Δ63 rad6∆::KanMX* |

**Table S3: Plasmids**

| Name/ Alias | Insert Description | *E. coli* Marker | *S. cerevisiae* Marker | Backbone | Type | Source/Ref |
|---|---|---|---|---|---|---|
| pRS314/ KB169 | Empty Vector | AmpR | *TRP1* | - | *CEN/ARS* | (28) |
| pQLinkN/ KB950 | pQLinkN link backbone | AmpR | - | - | Bacterial expression | (6) |
| KB1095 | 2μ plasmid of *E. coli* Tyr tRNA synthetase; *E. coli* tRNATyr amber suppressor tRNA. | AmpR | *LEU2* | pLH157 | 2μ | (8) |
| KB1167 | *RAD6-13xMyc* | AmpR | *TRP1* | pRS314 | *CEN/ARS* | This study |
| KB1228 | *3xHSV-RTF1* | KanR | *TRP1* | - | 2μ | (8) |
| KB1441 | *HMD74-184-F108BPA-7xHis* | AmpR | - | pQLinkN | Bacterial expression | This study |
| pEVOL-pBpF/ KB1444 | Aminoacyl-tRNA synthetase for BPA | CamR | - | - | Bacterial expression | (25) |
| KB1450 | *6xHis-MBP-TEV-V5-Rad6* | KanR | - | pLC3 | Bacterial expression | This study |
| pKA8/ KB1480 | Empty Vector | AmpR | - | - | Bacterial expression | Andrew VanDemark |
| KB1481 | *8xHis-TEV-V5-Rad6* | AmpR | - | pKA8 | Bacterial expression | This study |
| KB1491 | *8xHis-TEV-Rad6* | AmpR | - | pKA8 | Bacterial expression | This study |
| KB1499 | *rad6-K14A-13xMyc* | AmpR | *TRP1* | pRS314 | *CEN/ARS* | This study |
| KB1500 | *rad6-R11A-13xMyc* | AmpR | *TRP1* | pRS314 | *CEN/ARS* | This study |
| KB1501 | *rad6-D12A-13xMyc* | AmpR | *TRP1* | pRS314 | *CEN/ARS* | This study |
| KB1509 | *8xHis-TEV-rad6-F13A* | AmpR | - | pKA8 | Bacterial expression | This study |
| KB1514 | *rad6-F13BPA-13xMyc* | AmpR | *TRP1* | pRS424 | 2μ | This study |
| KB1517 | *8xHis-TEV-rad6-R11A* | AmpR | - | pKA8 | Bacterial expression | This study |
| KB1519 | *8xHis-TEV-rad6-M10A* | AmpR | - | pKA8 | Bacterial expression | This study |
| pUB23/ KB1526 | Ubiquitin-Arg-β-galactosidase reporter for testing the N-end rule pathway | AmpR | *URA3* | - | 2μ | Bachmair and Vershavsky, 1989 |
| KB1539 | *rad6-S2A-13xMyc* | AmpR | *TRP1* | pRS314 | *CEN/ARS* | This study |
| KB1540 | *rad6-T3A-13xMyc* | AmpR | *TRP1* | pRS314 | *CEN/ARS* | This study |

| | | | | | | |
|---|---|---|---|---|---|---|
| KB1541 | *rad6-P4A-13xMyc* | AmpR | *TRP1* | pRS314 | *CEN/ARS* | This study |
| KB1542 | *rad6-R6A-13xMyc* | AmpR | *TRP1* | pRS314 | *CEN/ARS* | This study |
| KB1543 | *rad6-R7A-13xMyc* | AmpR | *TRP1* | pRS314 | CEN/ARS | This study |
| KB1544 | *rad6-R8A-13xMyc* | AmpR | *TRP1* | pRS314 | *CEN/ARS* | This study |
| KB1545 | *rad6-L9A-13xMyc* | AmpR | *TRP1* | pRS314 | *CEN/ARS* | This study |
| KB1556 | *8xHis-TEV-rad6-S2A* | AmpR | - | pKA8 | Bacterial expression | This study |
| KB1557 | *8xHis-TEV-rad6-T3A* | AmpR | - | pKA8 | Bacterial expression | This study |
| KB1558 | *8xHis-TEV-rad6-R6A* | AmpR | - | pKA8 | Bacterial expression | This study |
| KB1559 | *8xHis-TEV-rad6-R8A* | AmpR | - | pKA8 | Bacterial expression | This study |
| KB1571 | *rad6-M10A-13xMyc* | AmpR | *TRP1* | pRS314 | *CEN/ARS* | This study |
| KB1572 | *rad6-F13A-13xMyc* | AmpR | *TRP1* | pRS314 | *CEN/ARS* | This study |
| KB1585 | *rad6-P4BPA-13xMyc* | AmpR | *TRP1* | pRS424 | 2µ | This study |
| KB1586 | *rad6-R7BPA-13xMyc* | AmpR | *TRP1* | pRS424 | 2µ | This study |
| KB1587 | *8xHis-TEV-rad6-L9A* | AmpR | - | pKA8 | Bacterial expression | This study |
| KB1588 | *8xHis-TEV-rad6-D12A* | AmpR | - | pKA8 | Bacterial expression | This study |
| KB1686 | *rad6-R6Q-13xMyc* | AmpR | *TRP1* | pRS314 | *CEN/ARS* | This study |
| KB1688 | *rad6-R6A,M10A-13xMyc* | AmpR | *TRP1* | pRS314 | *CEN/ARS* | This study |
| KB1689 | *8xHis-TEV-rad6-R6Q* | AmpR | - | pKA8 | Bacterial expression | This study |
| KB1690 | *8xHis-TEV-rad6-R6A,M10A* | AmpR | - | pKA8 | Bacterial expression | This study |
